# Hypothalamic Protein Profiling from Mice Subjected to Social Defeat Stress

**DOI:** 10.1101/2023.01.09.523315

**Authors:** Shiladitya Mitra, Ghantasala S Sameer Kumar, Anumita Samanta, Suman S Thakur

## Abstract

The Hypothalmic-pituitary axis also known as the HPA axis is central to stress response. It also acts as the relay centre between the body and the brain. We analysed hypothalamic proteome from mice subjected to chronic social defeat paradigm using iTRAQ based quantitative proteomics in identify changes associated with stress response. We identified greater than 2000 proteins processing our samples analysed through Q-Exactive (Thermo) and Orbitrap Velos (Thermo) at 5% FDR. Analysis of data procured from the runs showed that the proteins whose levels were affected belonged primarily to mitochondrial and metabolic processes, translation, complement pathway among others. We also found increased levels of fibrinogen, myelin basic protein (MBP) and neurofilaments (NEFL, NEFM, NEFH) in the hypothalamus from socially defeated mice. Interestingly these proteins are found upregulated in blood and CSF of subjects exposed to trauma and stress. Since hypothalamus is in direct contact of blood and CSF, their utility as biomarkers in depression holds an impressive probability and should be validated in clinical samples.

## INTRODUCTION

Depression is one of the major causes of morbidity worldwide. Depression can arise due to genetic as well as environmental factors. Despite a high rate of prevalence of depression, its causes and proper treatment still remains elusive (Hollis and Kabbaj, 2014). In this context, animal models of depression are of interest as they aid in understanding the etio-pathology of depression, finding molecular causes and well as diagnostic methods and screening of drugs. Social defeat model has been designed in a manner that it mimics multiple human situations like bullying, getting beaten up, physical abuse etc (Krishnan and Nestler, 2011). This make this model an interesting choice to be studied and analysed, results of which may be correlated to human beings.

Multiple brain regions have been studied that may have a profound influence on the pathology of depression. This includes the hippocampus, amygdala, nucleus accumbens, prefrontal cortex etc (Nestler *et al*, 2002). The hypothalamic pituitary axis (HPA) is at the foremost in response of the animal to stress as it acts as a relay centre for the brain and the body. Theoretically, modulation of the HPA axis can make an animal resilient or more susceptible to stress (Pfau and Russo, 2015). The expression and release of gluco-corticoids-which are key to stress response, are regulated by the HPA axis (Reul *et al*, 2015; Sapolsky *et al*, 2000). The hypothalamus is the main orchestrator of the HPA axis.

While the action of HPA axis when an animal is subjected to stress, is known, interestingly not much is known about other molecular changes in the hypothalamus that might be instrumental in understanding stress response - particularly in case when an animal is subjected to social defeat stress. Molecular changes that may happen to the hypothalamus when an animal is subjected to chronic stress may reveal the mode of action of the hypothalamic regions and may also indicate potential bio-markers. Mass spectrometry, particularly iTRAQ based quantitative proteomics has developed into a powerful tool over the past decade to identify potential bio-marker(s) (Crutchfield *et al*, 2016; DeSouza *et al*, 2013; Street and Dear, 2010). Hence by employing proteomics to analyse hypothalamus of a mice subjected to social defeat stress, we attempted to find molecular changes that may be extrapolated to depressed human subjects.

## MATERIALS AND METHODS

### a) Animals used

We used C57Bl/6J mice at 3-4 months of age for our experiment. The mice were housed in our in-house facility with 12 hours of light and dark cycle. Food was provided ad-libitum. CD1 mice were chosen as the aggressor strain. For this purpose CD1 mice, which were retired breeders, were used. All the experiments were approved by Institutional Animal Ethics Committee (IAEC, CCMB; Reg. No. CPCSEA 20/1999).

### b) Social Defeat Experiment

We followed a modified protocol that has been previously described (Golden *et al*, 2011; Wilkinson *et al*, 2009). Briefly, CD1 and C57Bl/6J mice were housed together with a transparent partition between them. Once a day for 10 minutes over 10 days, C57Bl/6J mice were placed in the partition of the CD1 mice (n=10) and were allowed to bully the former. Care was taken that no physical injury was inflicted on the C57Bl/6J strain. A Latin square approach was followed to prevent habituation. A separate set of C57Bl/6J mice (n=7) were not exposed to social defeat served as wild-type controls.

### c) Behavioral Experiment

We performed social avoidance experiment as a test for defeat. The protocol followed has been discussed in other reports (Golden *et al*, 2011; Wilkinson *et al*, 2009). In short, C57Bl/6J mice were kept in an open field with a transparent grilled enclosure/cage at one side. It was either empty or contained a CD1 aggressor mice. Area around the box was demarcated as interaction zone. Using Noldus Ethovision we tracked the duration and the path of the C57Bl/6J mice in the interaction zone in the presence or absence of the aggressor. A ratio between the two was calculated as the interaction ratio and was measured as a percentage of total time spent in both conditions.

### d) Proteomic Experiment

Samples were processed and analysis was carried out using methods described previously(Mitra *et al*, 2018). Data are available via ProteomeXchange with identifier PXD005644. Briefly,

#### iTRAQ 4 plex

Three each of wild-type and social defeat mice were selected and hypothalamus were collected and pooled. We used 0.5% SDS to extract the proteins. After quantification using BCA kit (Thermo Fischer), we used 200ug of protein from both groups to proceed with labelling reaction. The protocol followed was as described by the manufacturer (ABSciex). TCEP (tris (2-carboxyethyl) phosphine) was used to reduce and methyl methanethiosulfonate (MMTS)- a cysteine blocker for reducing the samples. Samples were subsequently digested with trypsin (Promega Cat#:V511A) overnight using 1:20 (w/w) at 37°C. 100 ug of each sample was used for one technical replicate and labelled with iTRAQ 4plex (ABSciex) reagents as per manufacturer’s protocol. Peptides from wild type samples were labelled with 114 and 115 tags, while those from social defeat samples were labelled with 116 and 117 tags. After labelling, samples were pooled, desalted using C^18^ spin columns (Pierce®) as per manufacturer’s protocol. The samples were subsequently processed for LC-MS/MS analysis after SCX fractionation using protocol described in a previous study (Venugopal *et al*, 2013).

#### LC-MS/MS analysis

Samples were analyzed on UPLC (Dionex The UltiMate® 3000 HPLC) interfaced Orbitrap Analyser (Thermo Scientific, Bremen, Germany) or Q-Exactive. Labelled peptides were introduced in a 15 cm long column (EASY-Spray column ES800, 15 cm x 75 μm ID, PepMap C18, 3 μm) and heated to 30° C. Peptides were separated using linear gradient from 2% to 98% of buffer B (95 % acetonitrile and 0.1% formic acid) at a flow rate of 300 nl/min. The gradient length was 50 minutes. Normalized collision energy (NCE) was set to 27 for fragmentation. Precursor ions were selected based on the charge state with priority of 2+, 3+ and >3+. The dynamic exclusion was set as 30 seconds during data acquisition. The nano source was operated with 2.2 KV and the capillary temperature at 300°C.

#### Data analysis

The data was analysed using Proteome Discoverer (Thermo Scientific, Bremen, Germany) software. NCBI fasta (for mouse) database was used to search for peptides. The workflow consisted of spectrum files, spectrum selector, reporter ion quantification and Sequest. Parameters included Methylthio (C), iTRAQ label at N-terminus of peptide. Lysine were used as a fixed modifications, whereas oxidation of methionine (M) and deamination (N/Q) were used as variable modification. The parent and fragment mass error tolerance were set as 20 ppm and 0.2 Da respectively. Using data from runs from both the machines, we acquired a total 77,751 MS/MS spectra and a total of 23,100 sequences were searched. We applied 5% FDR in our analysis and proteins with minimum of 1 unique peptide were considered.

### e) Real Time analysis

cDNA was prepared from the hypothalamus of wild-type and social defeated C57Bl/6J mice (n=4) using protocol listed elsewhere (Mitra *et al*, 2016). Real time analysis was performed using Sybr Green (Invitrogen) reagent in ABI HT 4900 machine according the manufacturer’s instructions and with 58°C as the annealing temperature. The primers used for the experiment are listed below.

### f) Statistical analysis

Behavioral data were subjected to students t-test. Data from real time PCR were subjected to non-parametric test. Proteomics data was subjected to FDR analysis.

## RESULTS

The HPA axis is one of the major circuits that play an important role in an organism’s stress response. The hypothalamus is central to the HPA axis’ response to stress (Smith and Vale, 2006). We carried out 4plex iTRAQ quantitative proteomic experiment to study the molecular changes in the hypothalamus when mice were subjected to chronic stress specifically social defeat stress.

A total of 17 mice were used for this study. While 7 mice were used as unstressed controls, 10 mice were subjected to social defeat paradigm. Mice subjected to social defeat stress spent significantly less time (p<0.05) in the interaction zone with the presence of the target than without (Fig 1C). The non-stressed controls did not show any significant differences. Equivalently, the group of mice subjected to social defeat stress showed significantly lesser interaction ratio (p<0.05) as compared to the controls (Fig 1D).

**Figure 1.**
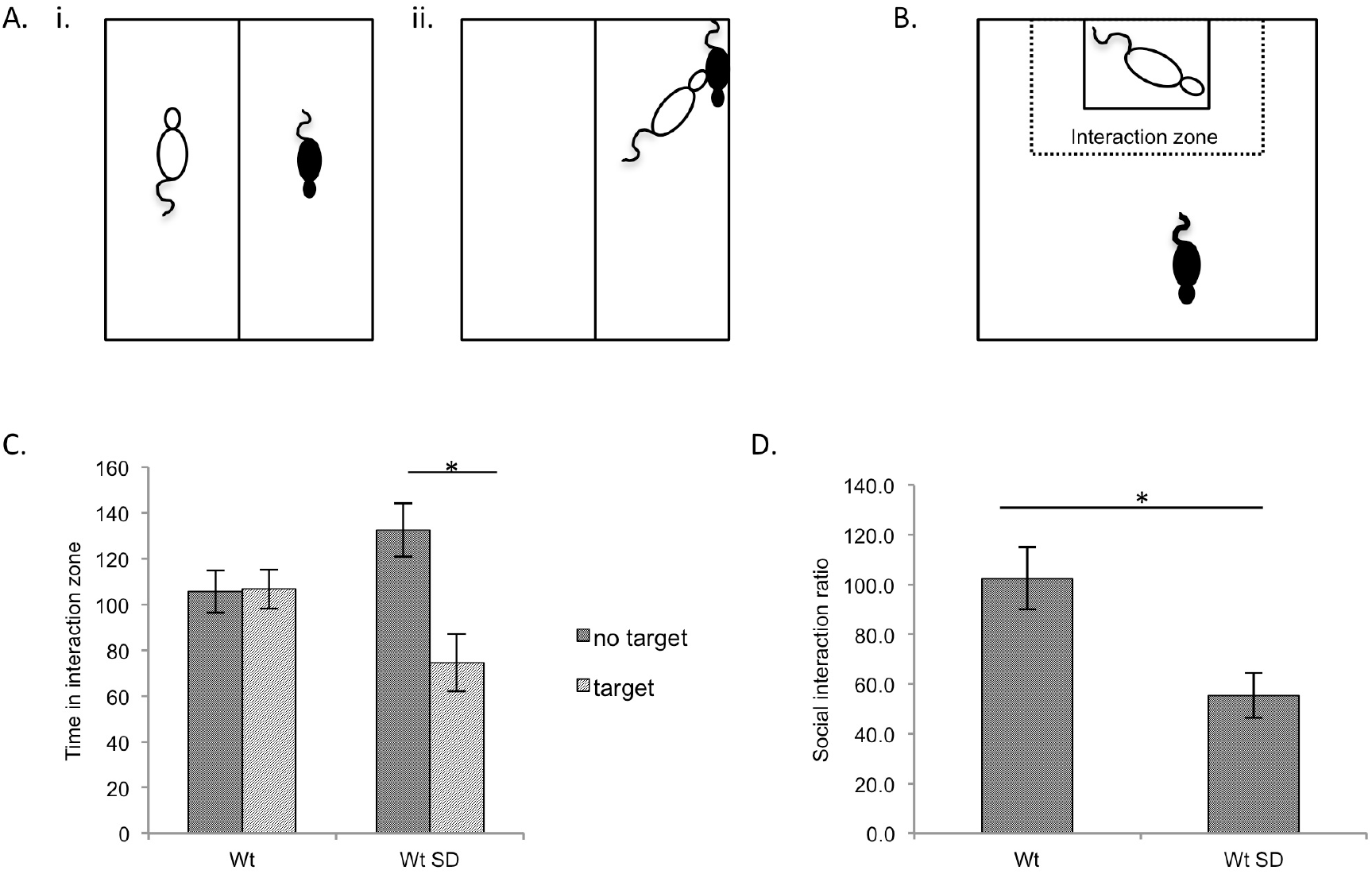
A. Social Defeat Stress procedure. (i) Male CD1 and C57Bl/6J are housed in one cage with transparent perforated partition in between. (ii) CD1 mice allowed to interact (bully) with the C57Bl/6J mice for 10 minutes every day for 10 days. B. Analysis of chronic stress. C57BL/6J mice of defeated and control groups are allowed to explore a box containing a target (CD1 mice) in one box. The social interaction ratio is calculated as the ratio between time spent in the interaction zone with the target compared to without the presence of the target. C. The time spent by the defeated mice (Wt SD) in the interaction zone was significantly lesser (p<0.05) in the presence of a target as compared to without the target. There was significant difference in the time spent in the interaction zone by the unstressed controls (Wt). D. The social interaction ratio was significantly lesser (p<0.05) in the defeated group (Wt SD) than the unstressed controls (Wt).

We used iTRAQ based quantitative proteomics to search for differentially expressed proteins in the hypothalamus of mice after chronic stress. In our analysis, combining runs from both Orbitrap Velos and Q-Exactive, we were able to identify more than 2000 unique proteins (Supplementary Table 1). We found a total of 181 proteins upregulated (>1.5 folds (116+117/113+114 or social defeat/unstressed) in the hypothalamus of the mice subjected to social defeat stress as compared to the unstressed controls (Supplementary Table 2). In our analysis we failed to find any significantly downregulated proteins. The most plausible reason behind this would be technical limitations of our runs. However, it has been reported that after social defeat, the number of genes that were downregulated in the hypothalamus were considerably lesser than those upregulated (Smagin *et al*, 2016).

The upregulated proteins were subjected to network analysis using STRING software. They were arranged based on KEGG and GO Biological Processes pathway analysis using the same program. The top 10 pathways have been listed in Table 1 and Table 2. The full list for altered Biological Pathways have been listed in Supplementary Table 3. Network analysis was carried out with medium confidence setting with only primary nodes. The proteins were clustered in 10 prominent interacting groups (Fig 2, Supplementary Table 4). The most prominent groups consisted of the proteins associated with translation (cluster 1), complement and coagulation pathway (cluster9), cell adhesion and myelination (cluster 8), cytoskeletal organisation and neurofilament (cluster 2), chromatin assembly and organisation (cluster 7) and mitochondrial and metabolic processes (cluster 4 and cluster 6).

**Table 1:**
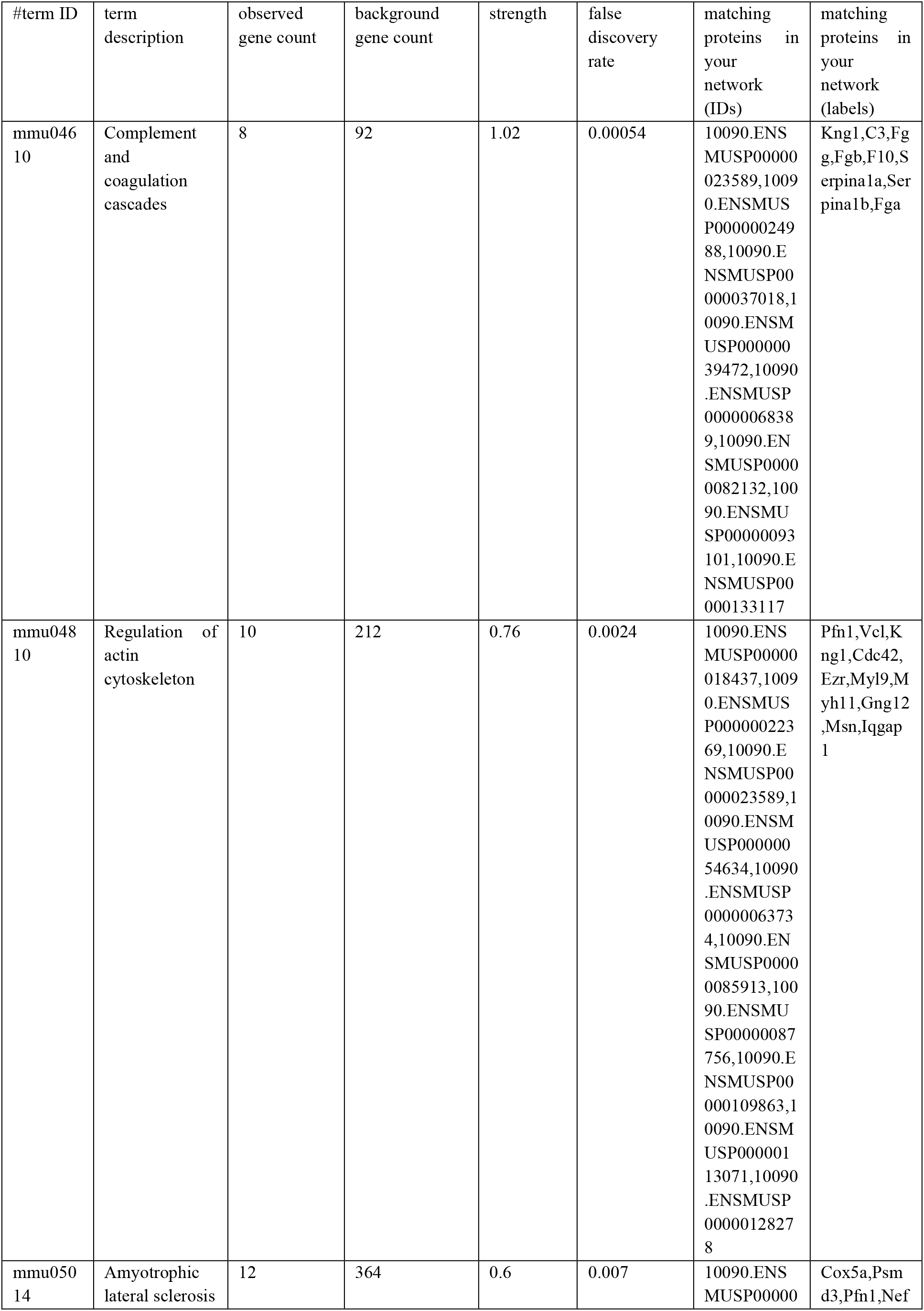
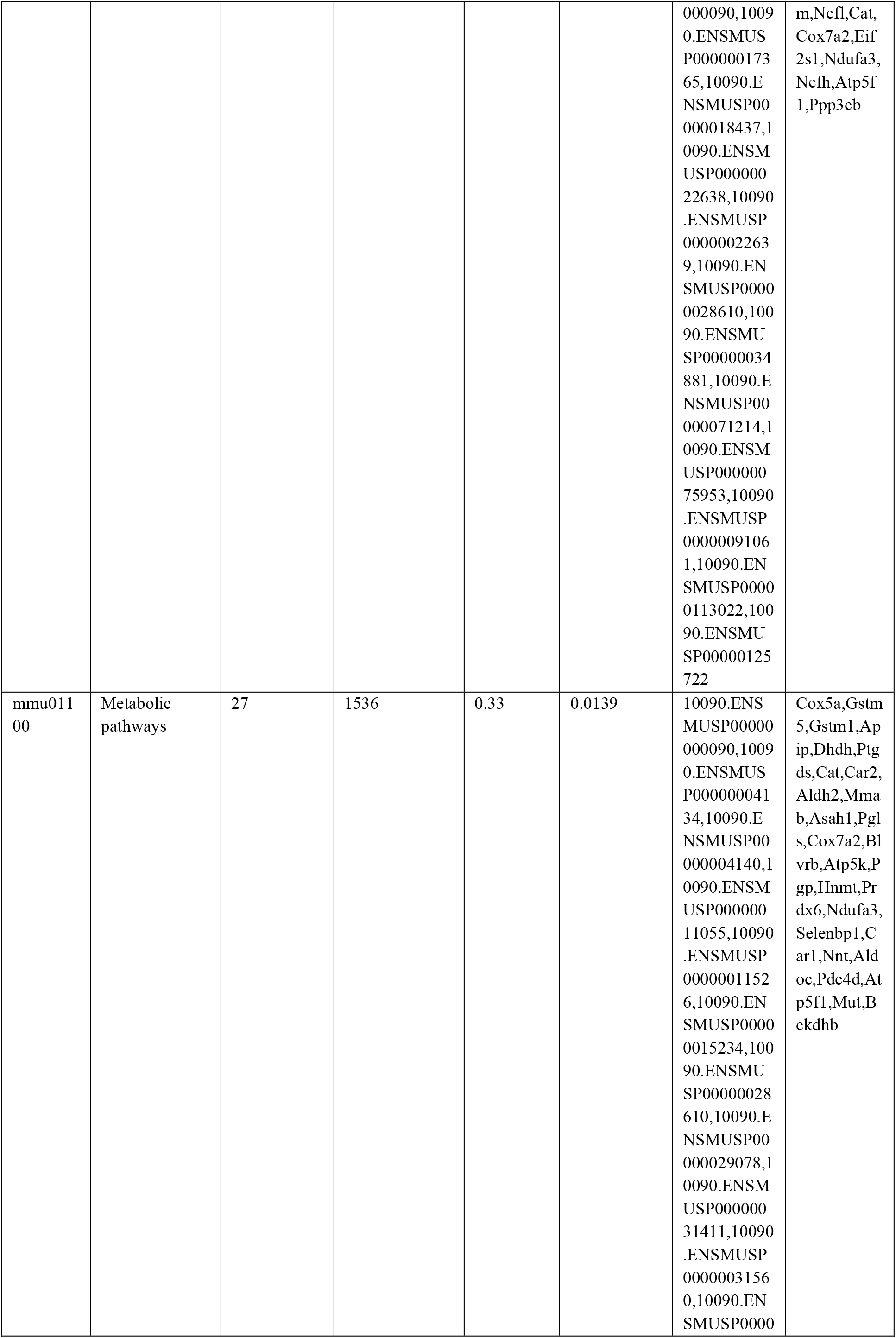
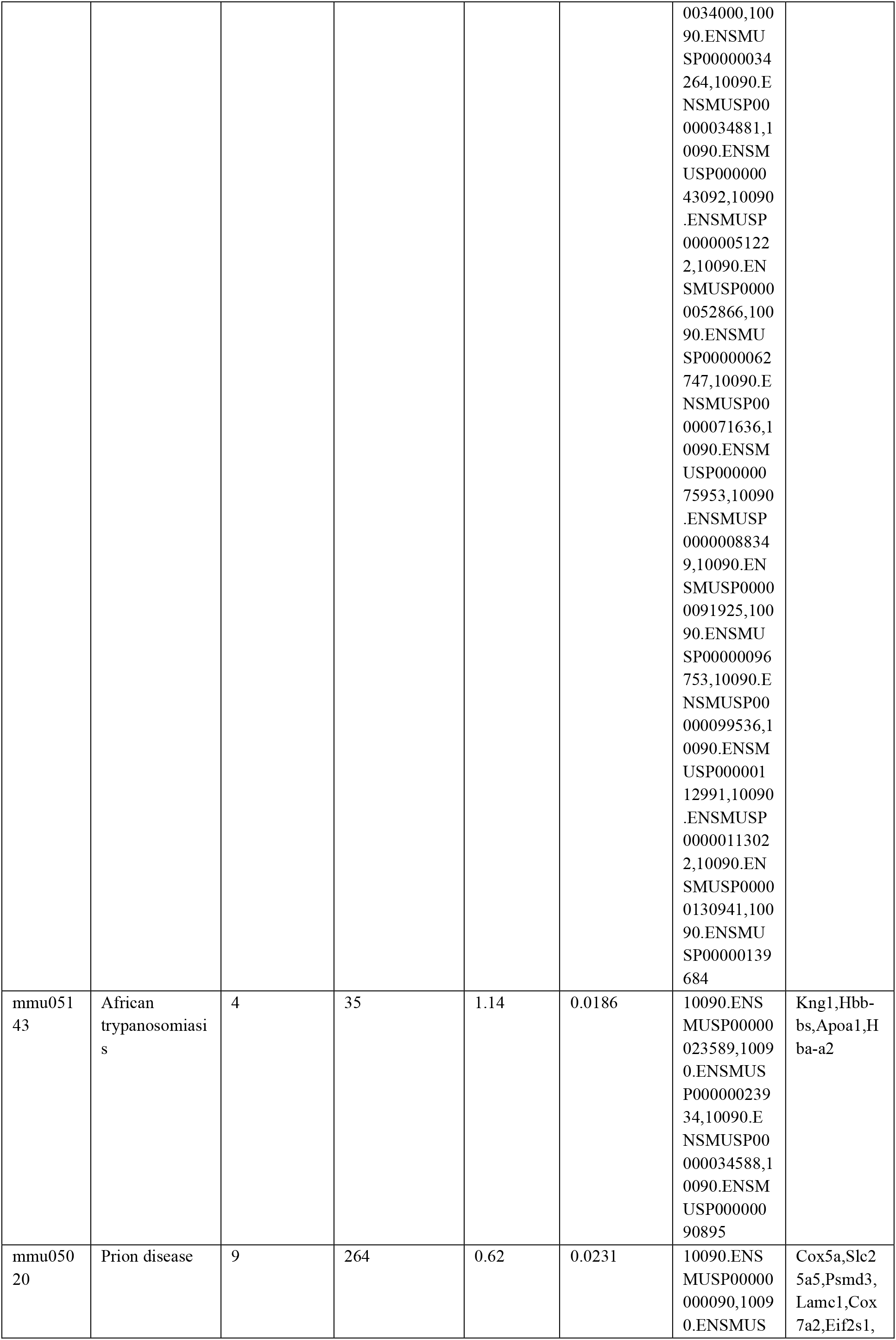
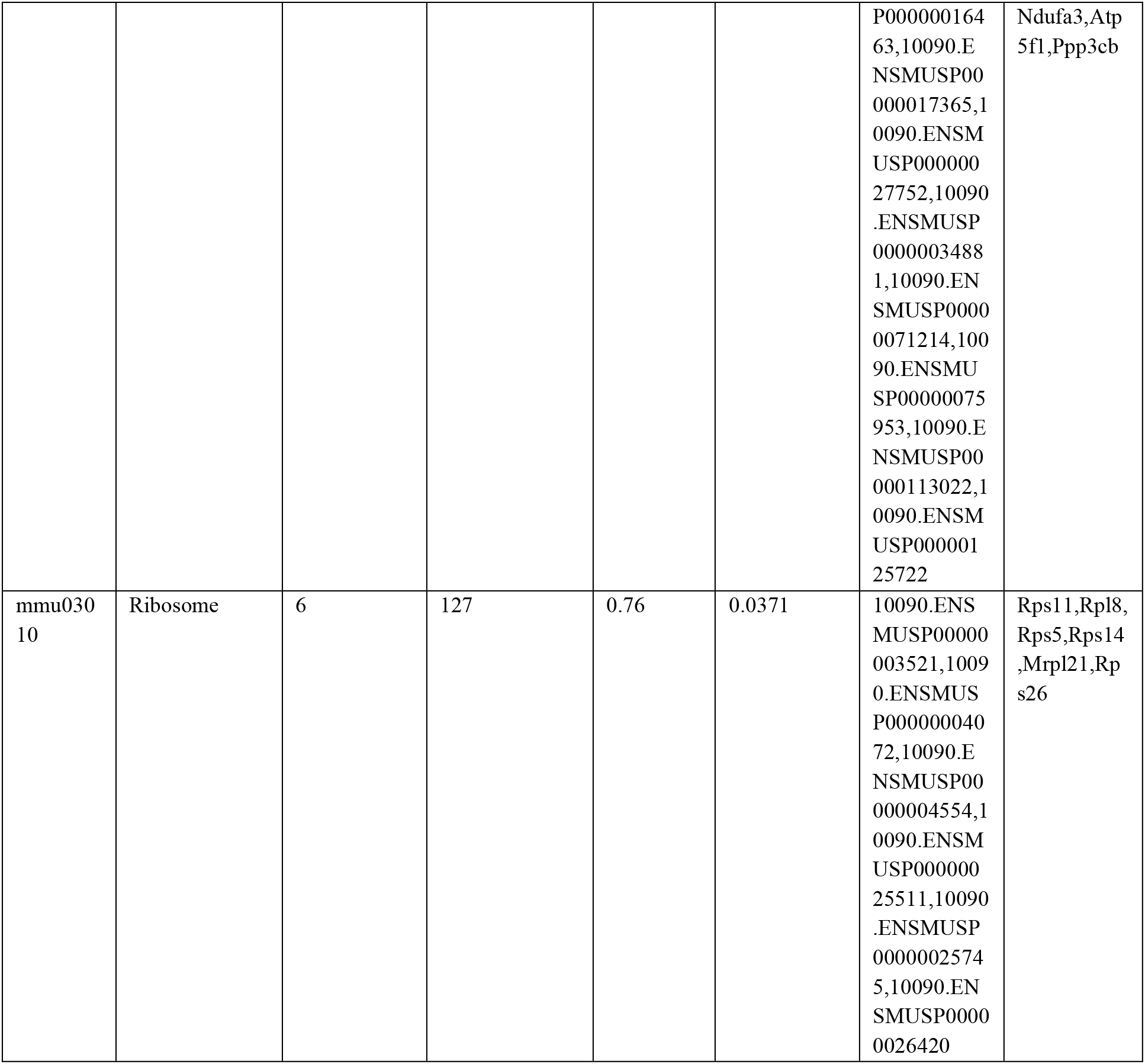
KEGG pathway analysis reveal the top 10 pathways affected

**Table 2 :**
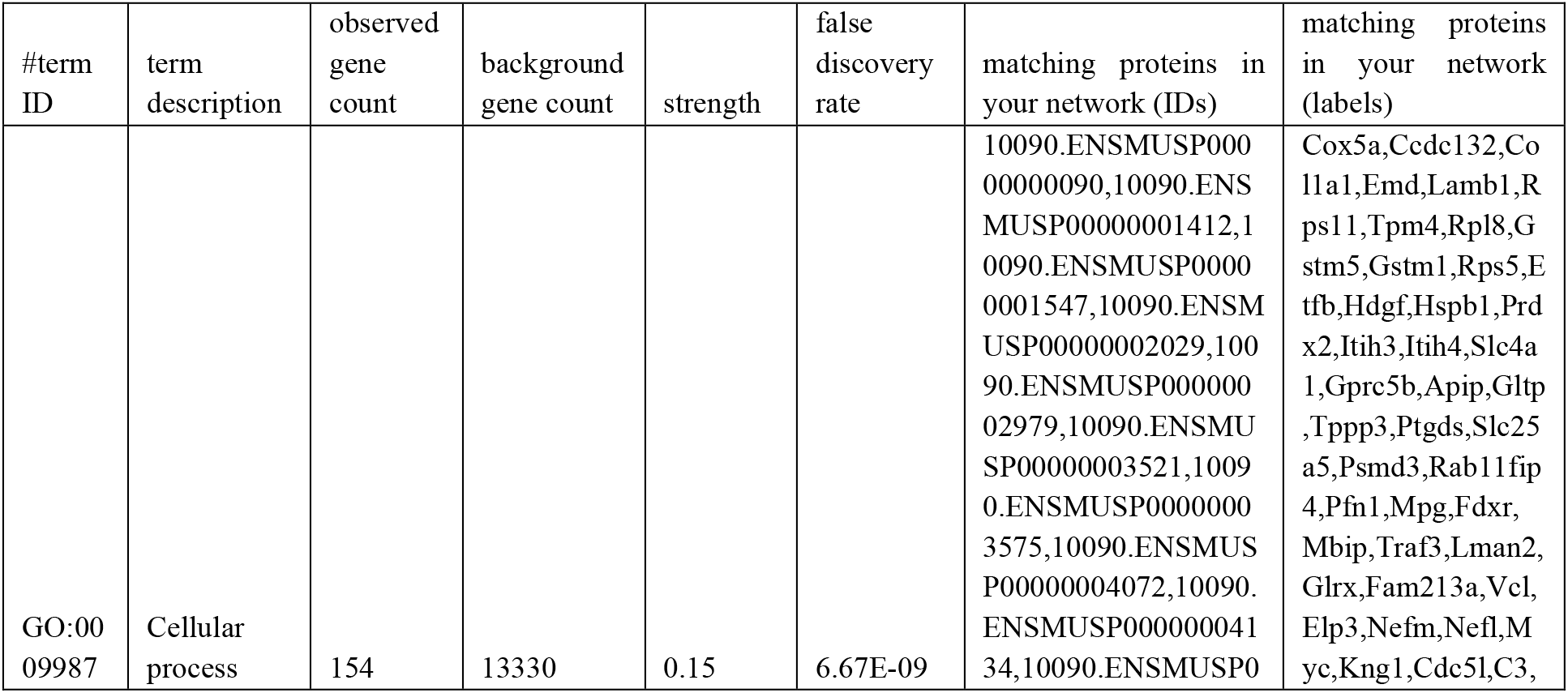
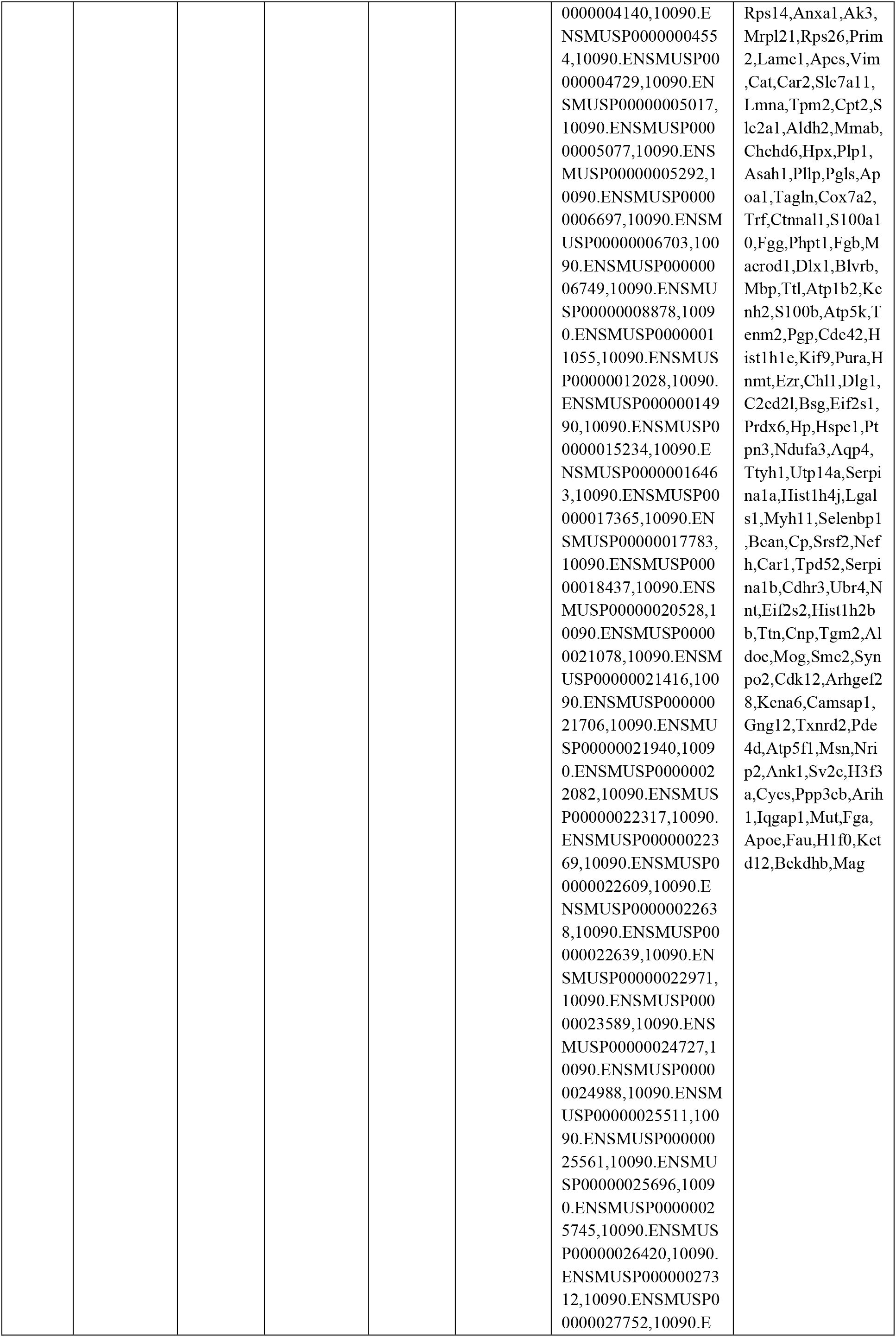
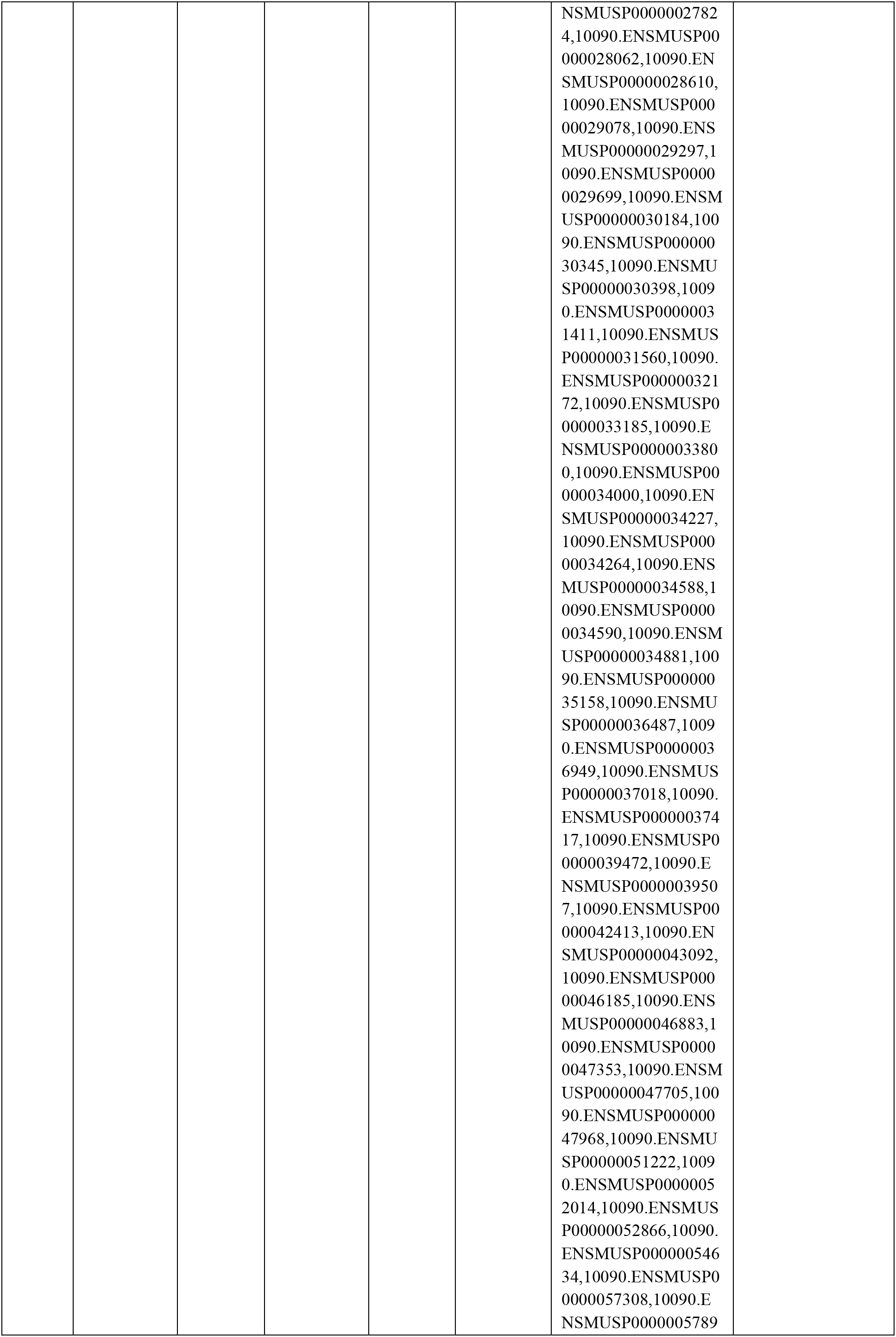
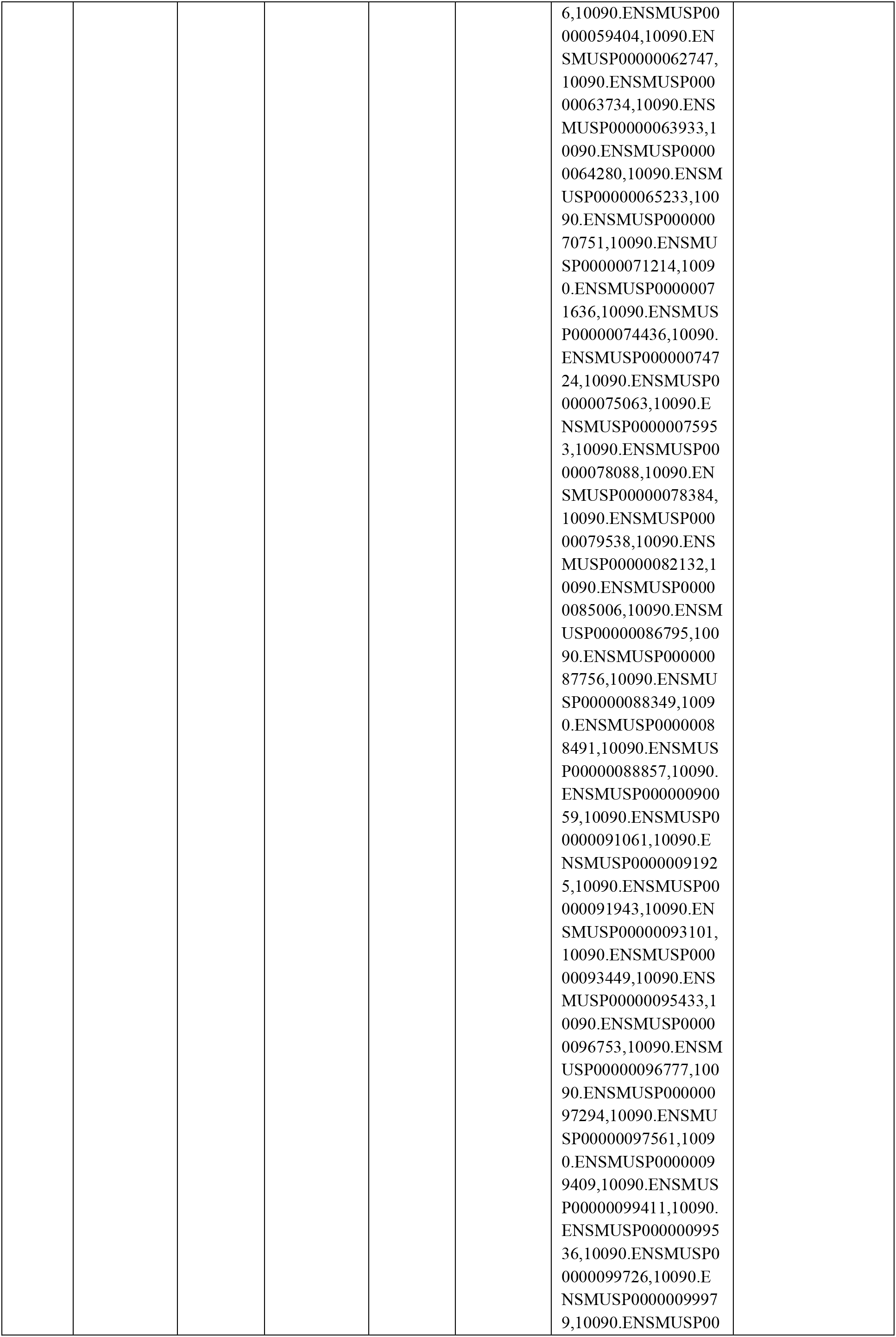
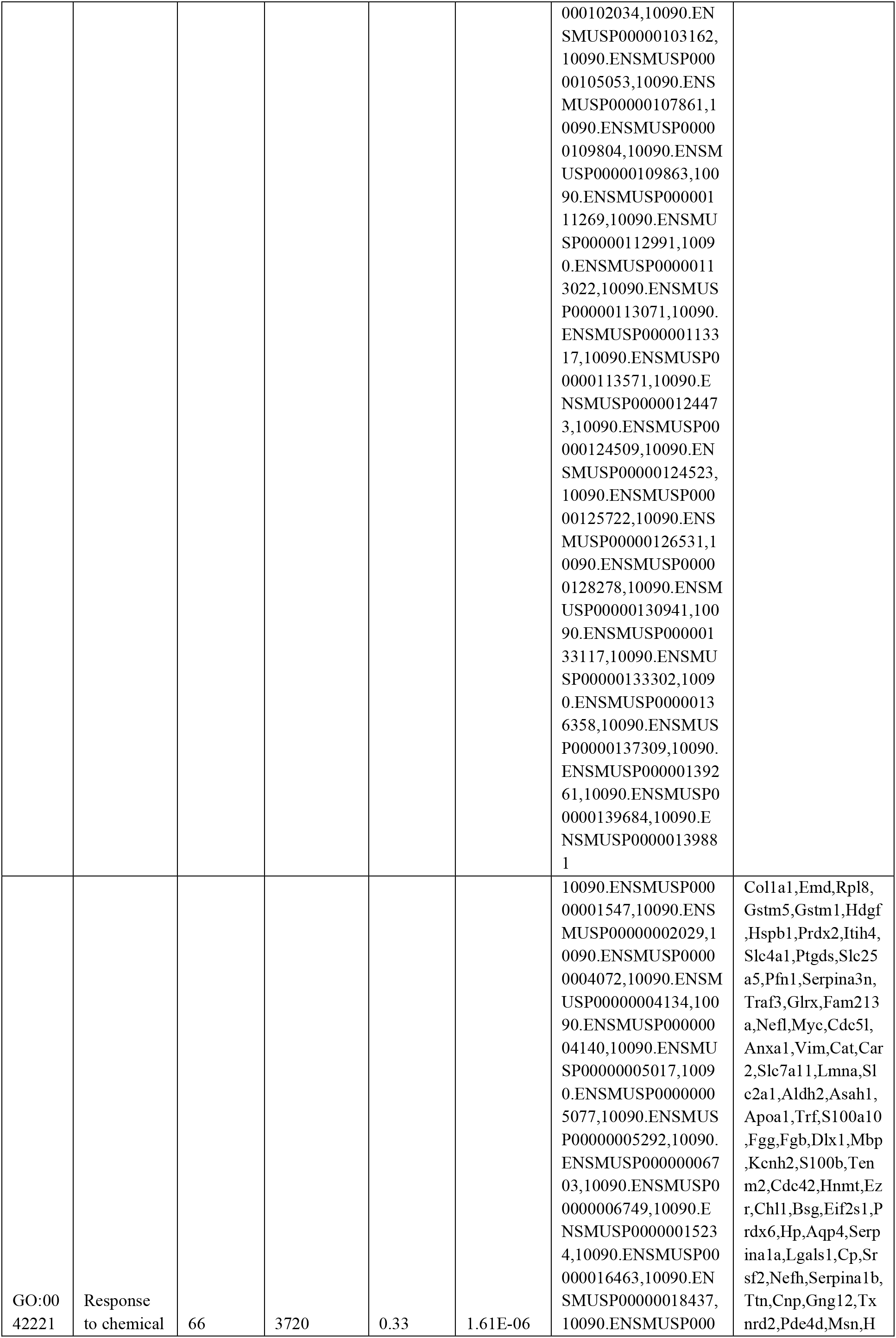
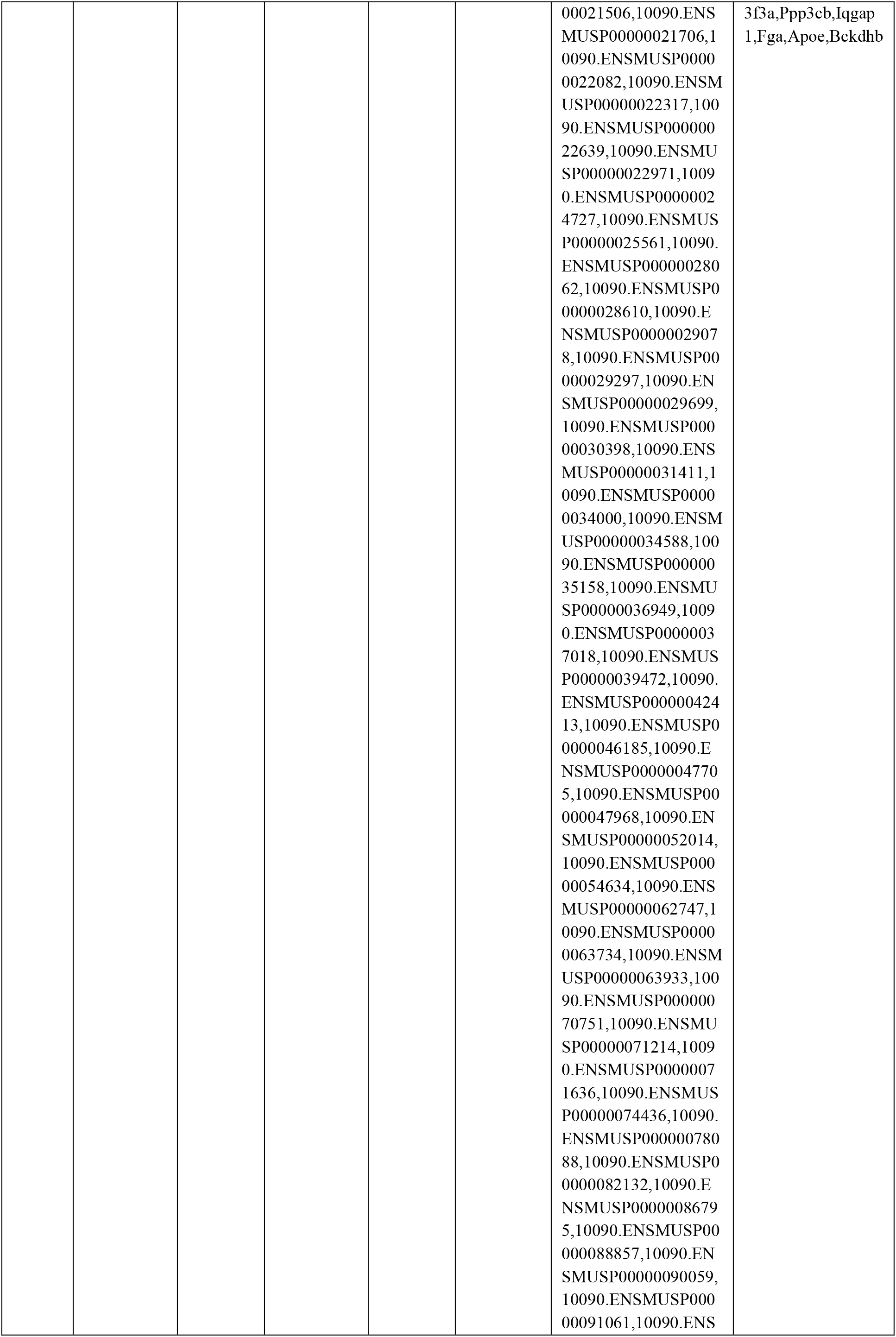
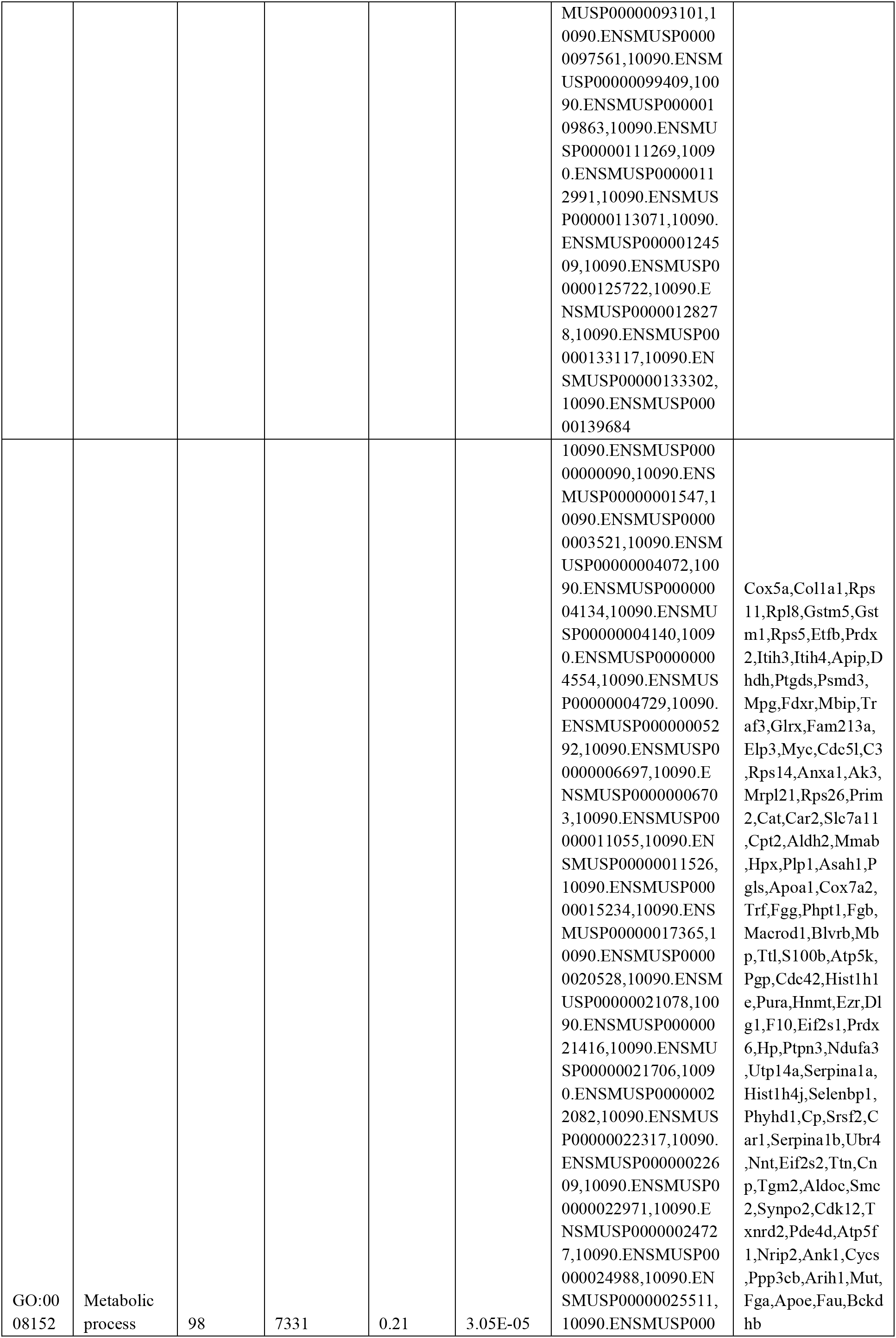
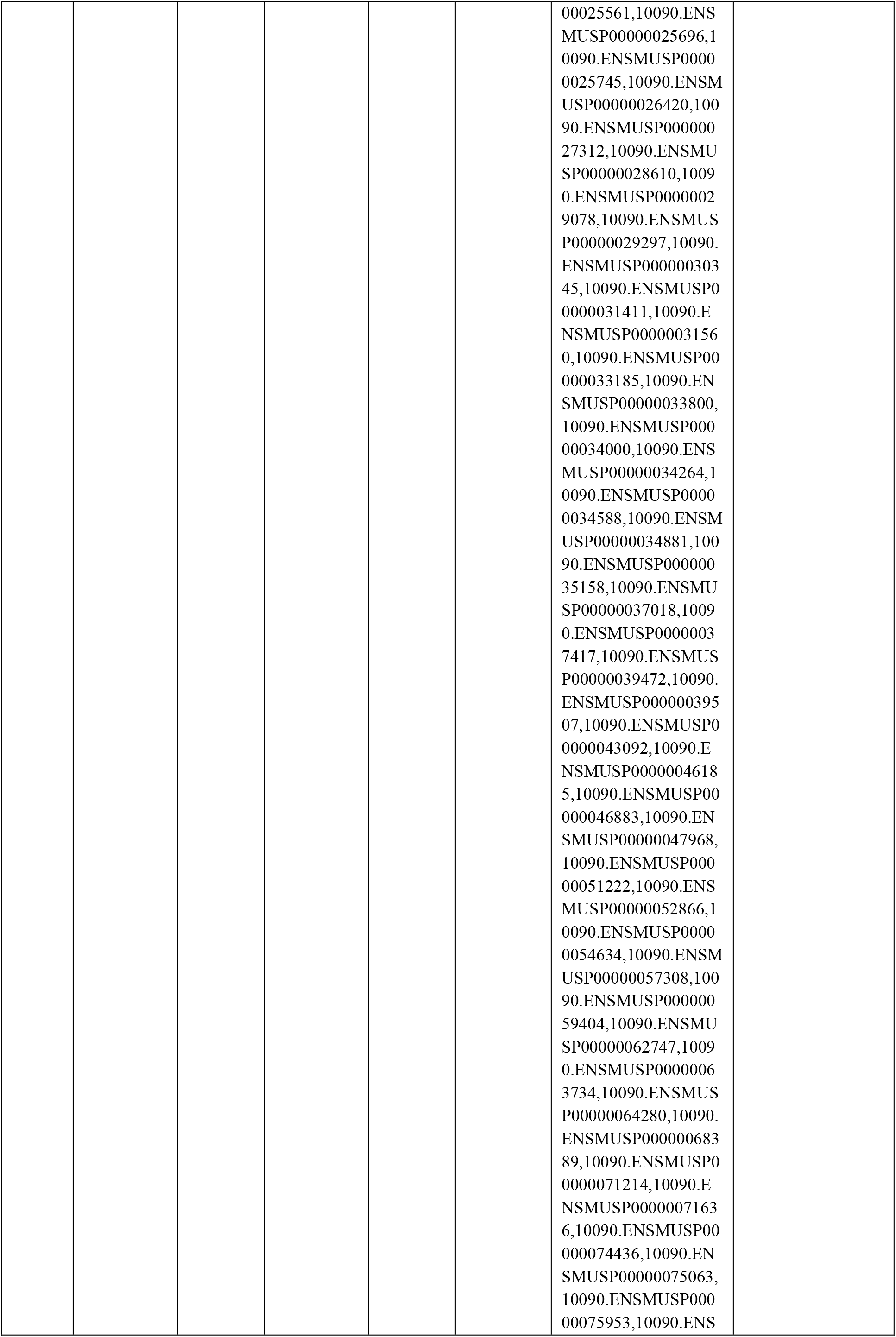
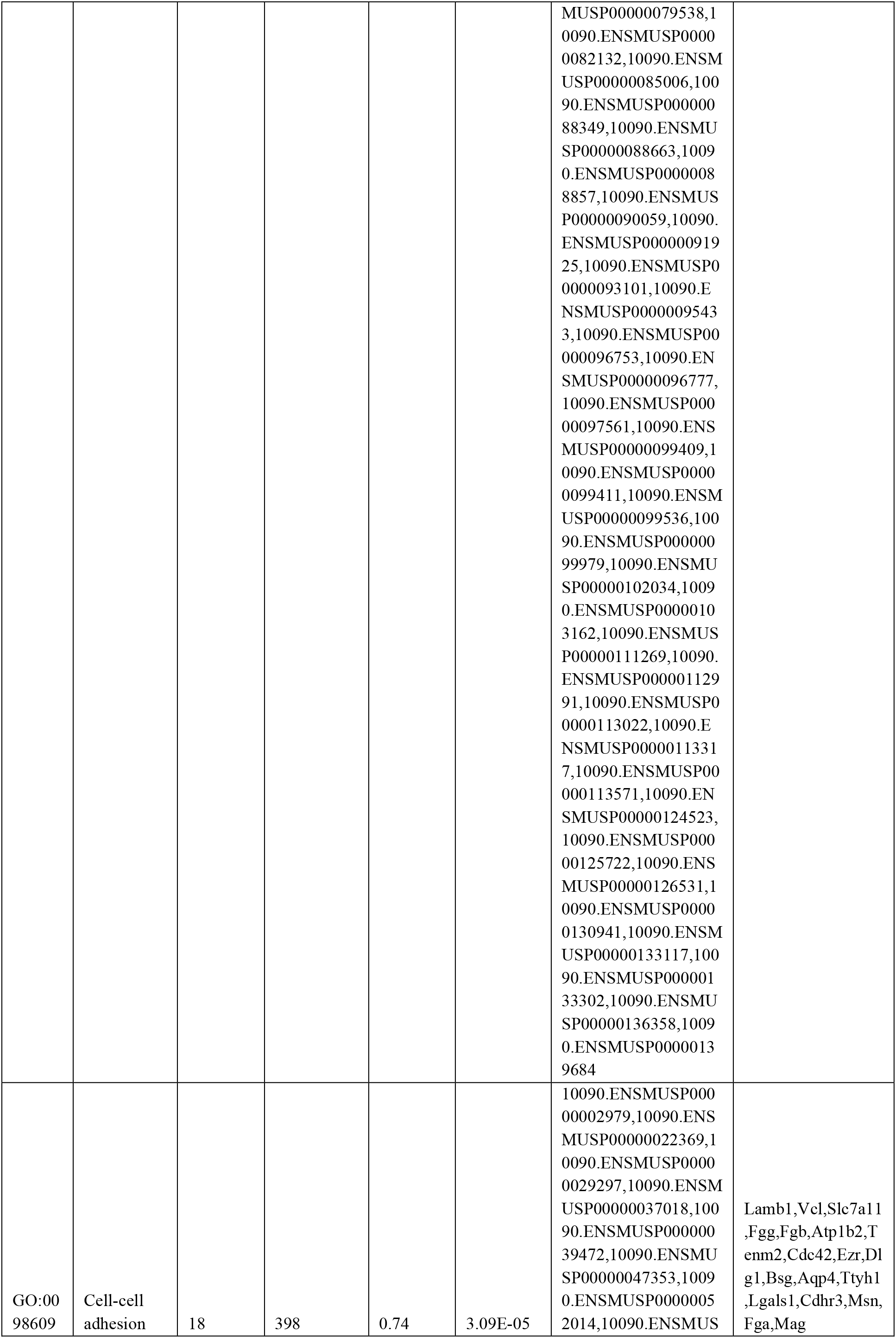
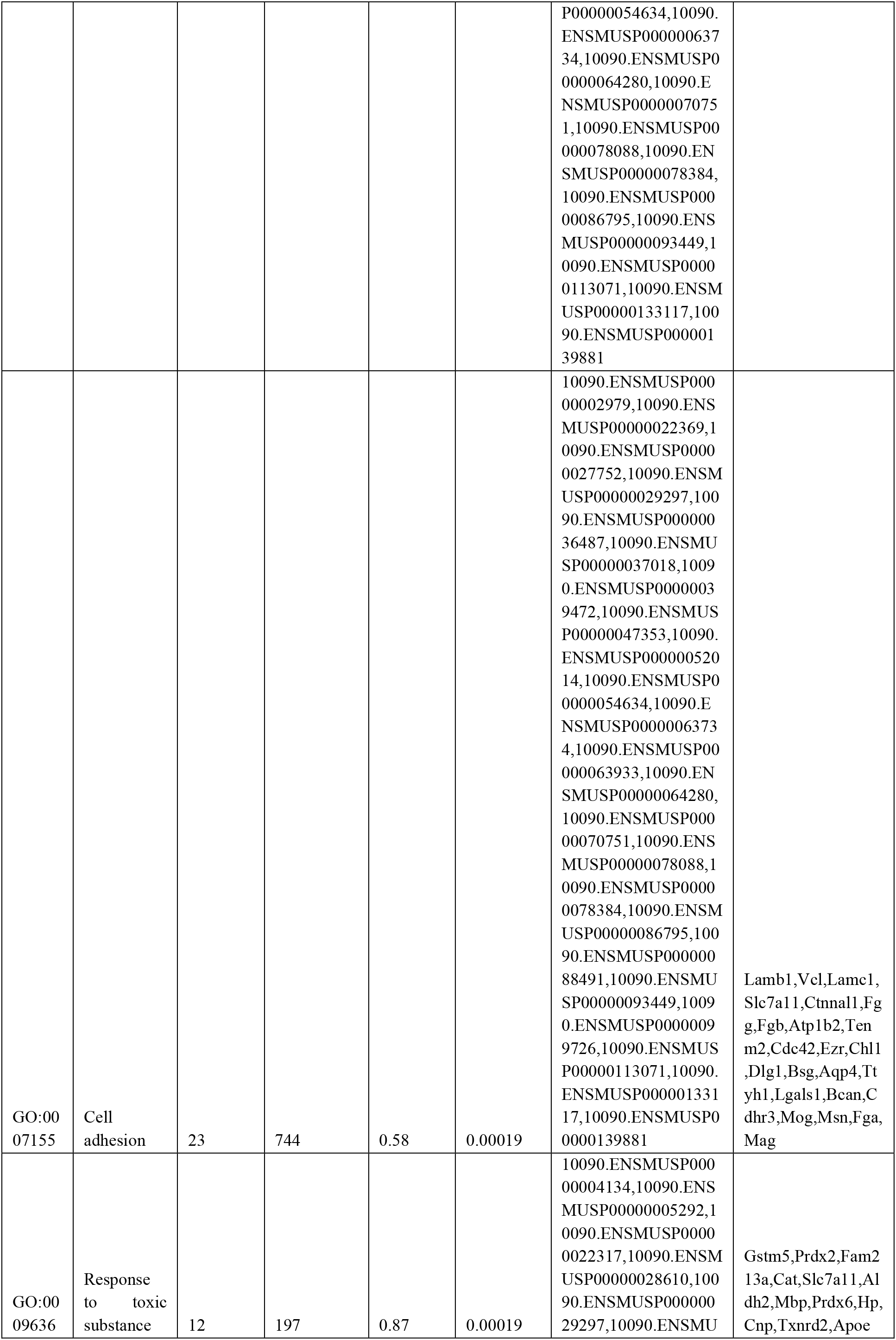
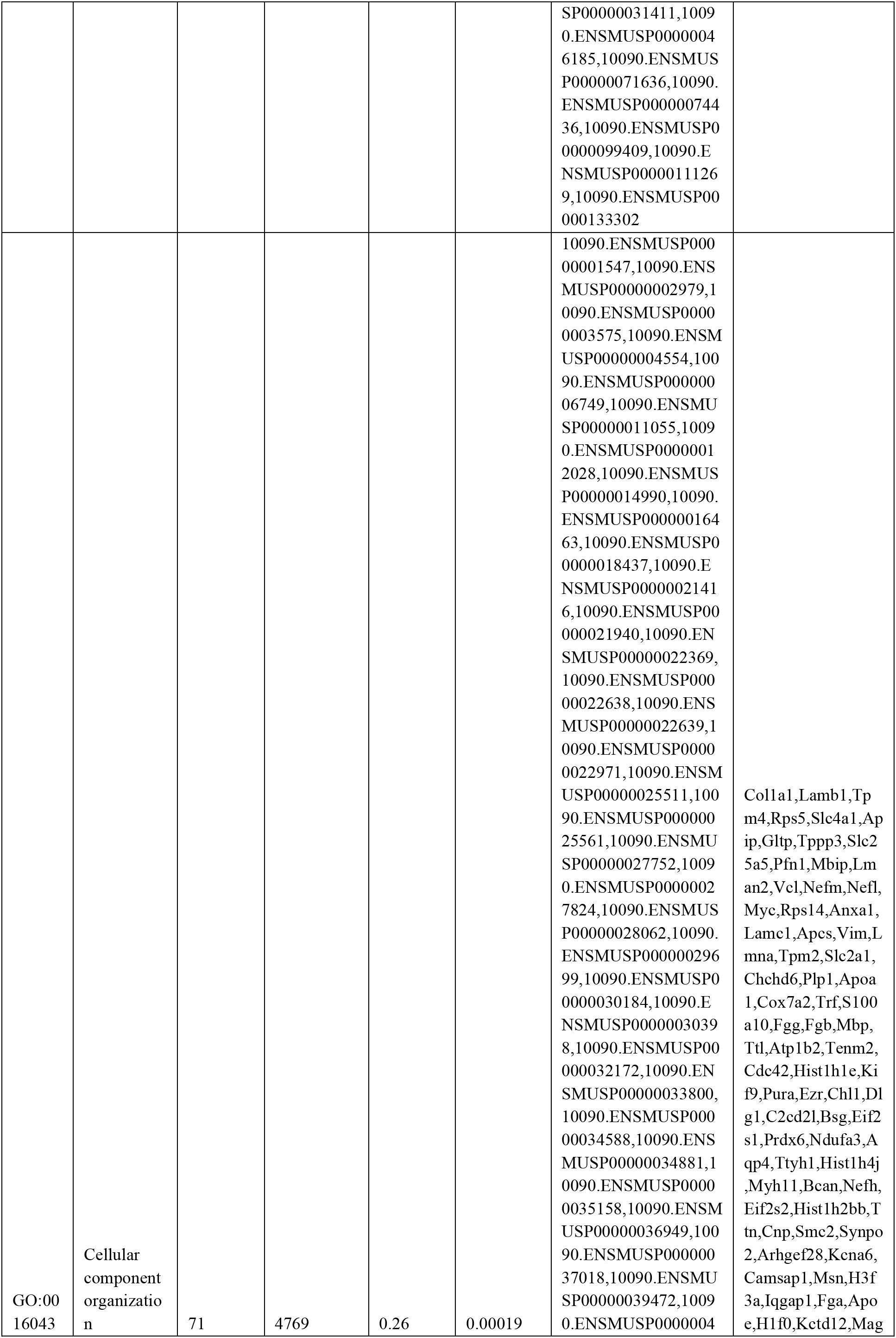
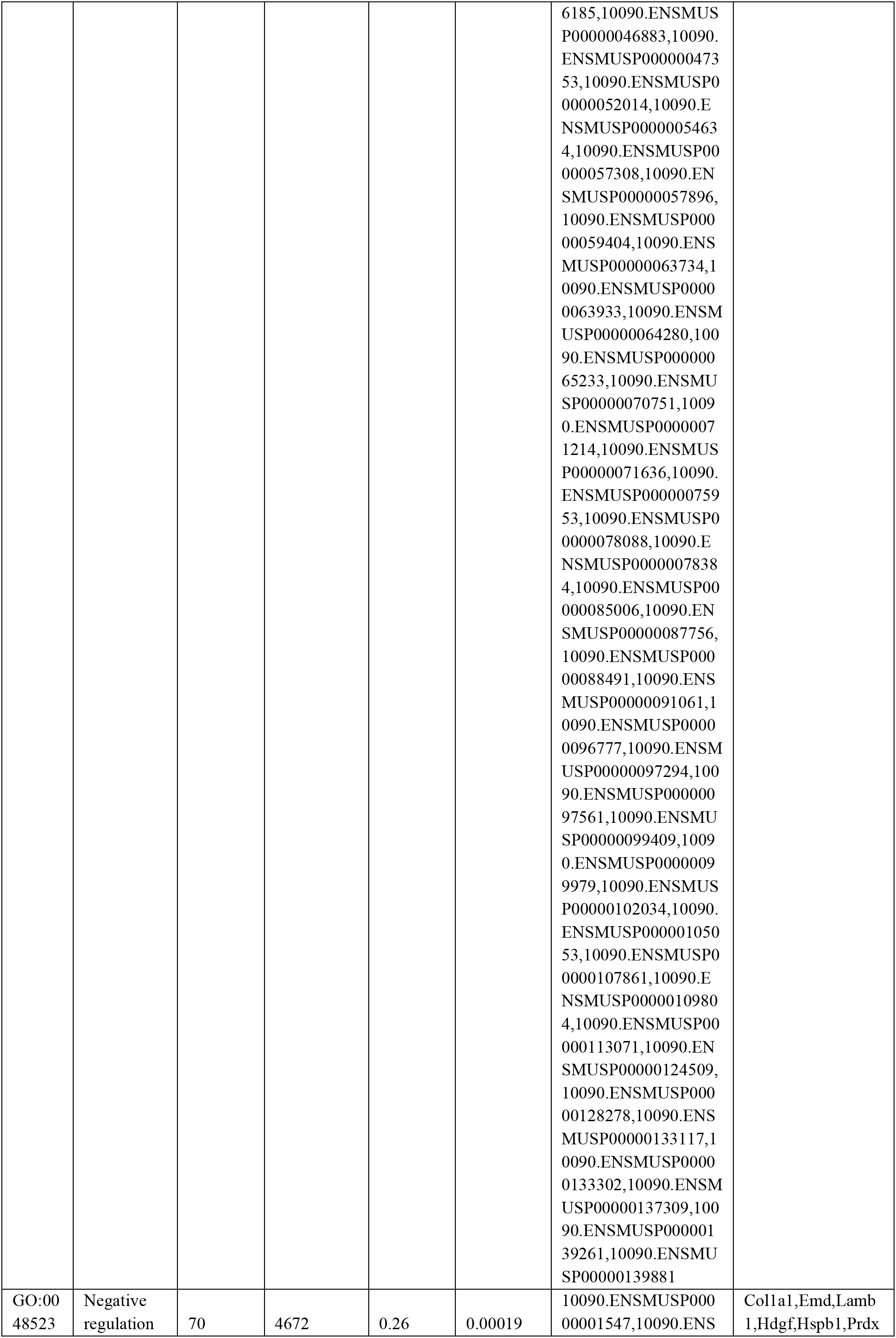
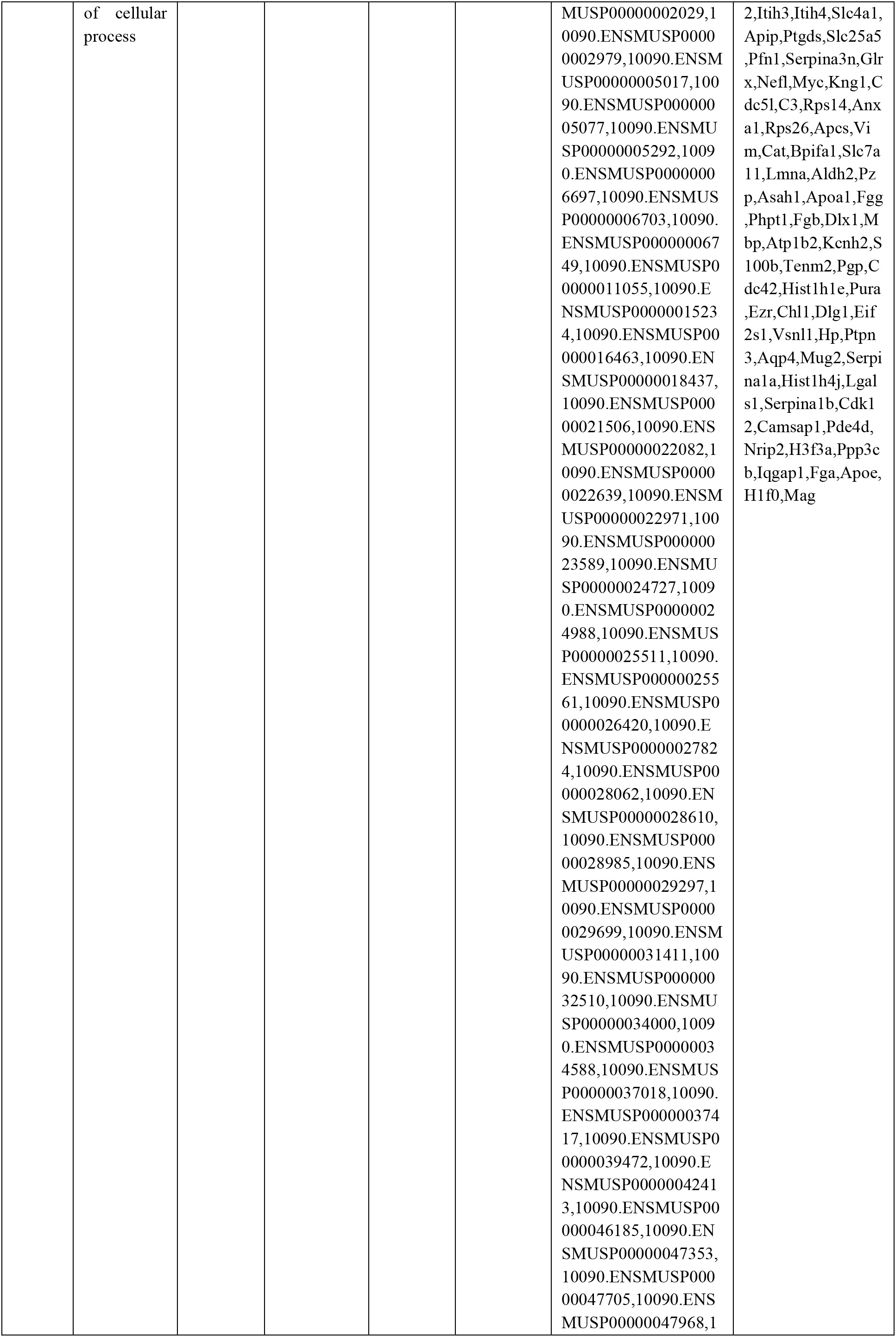
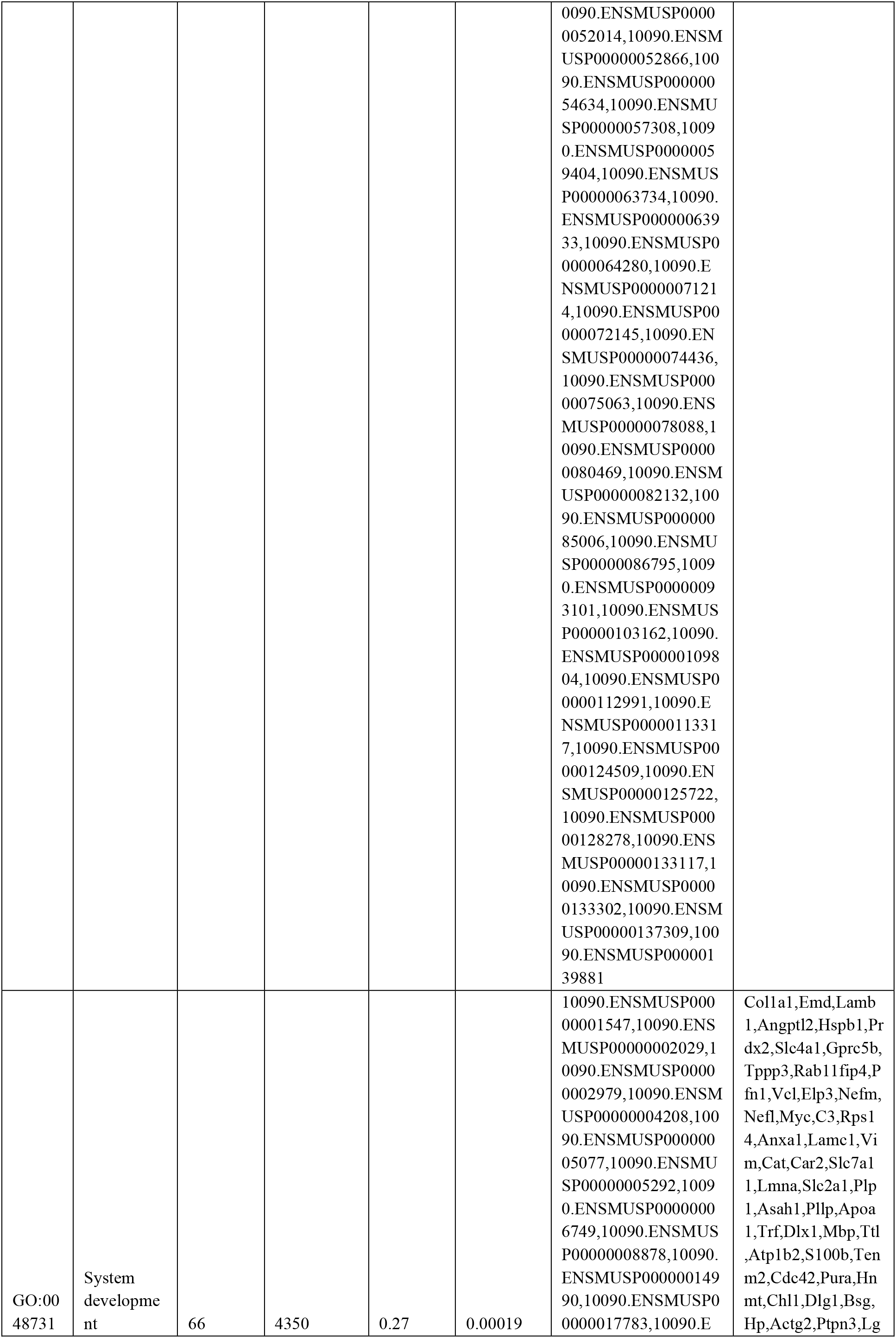
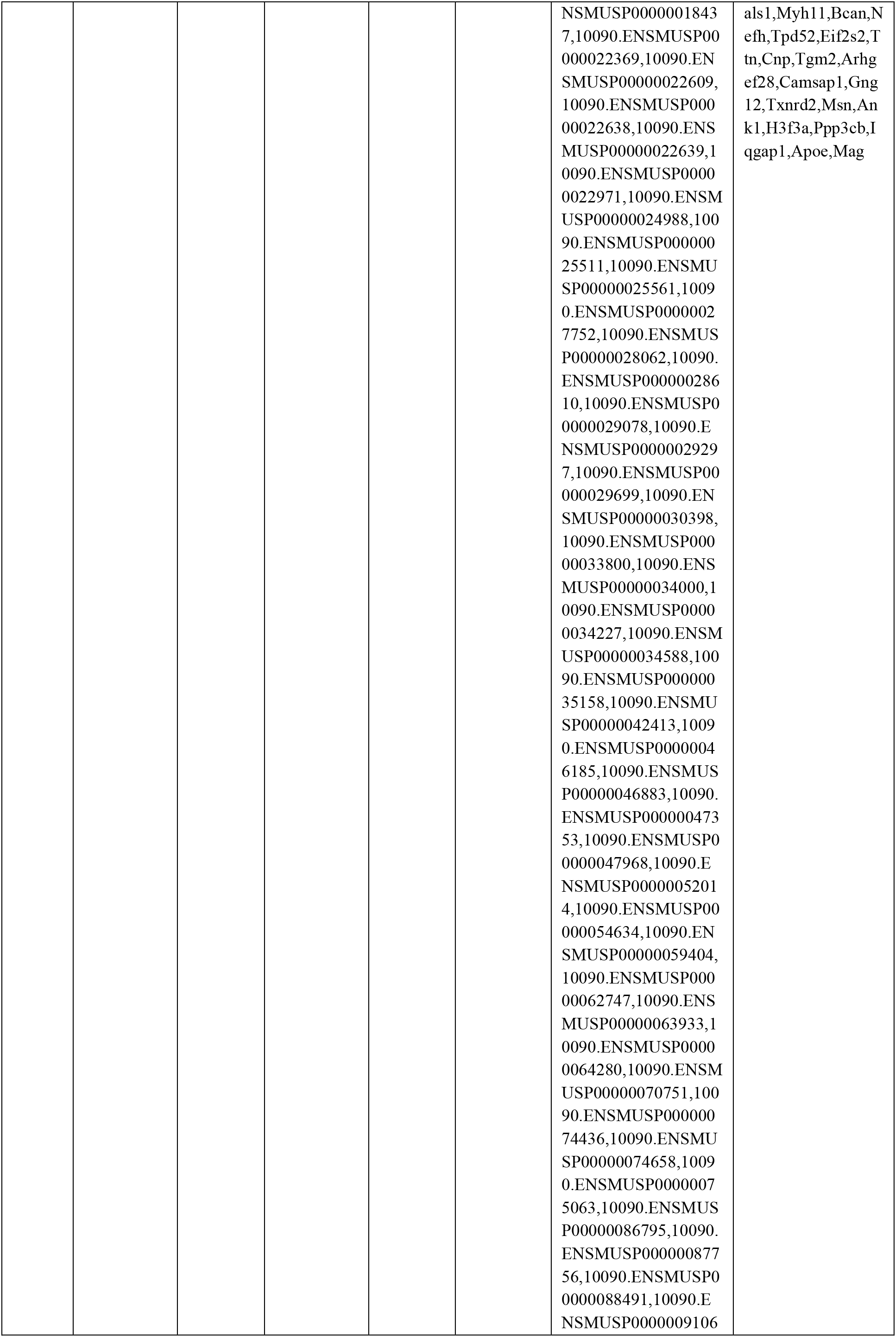
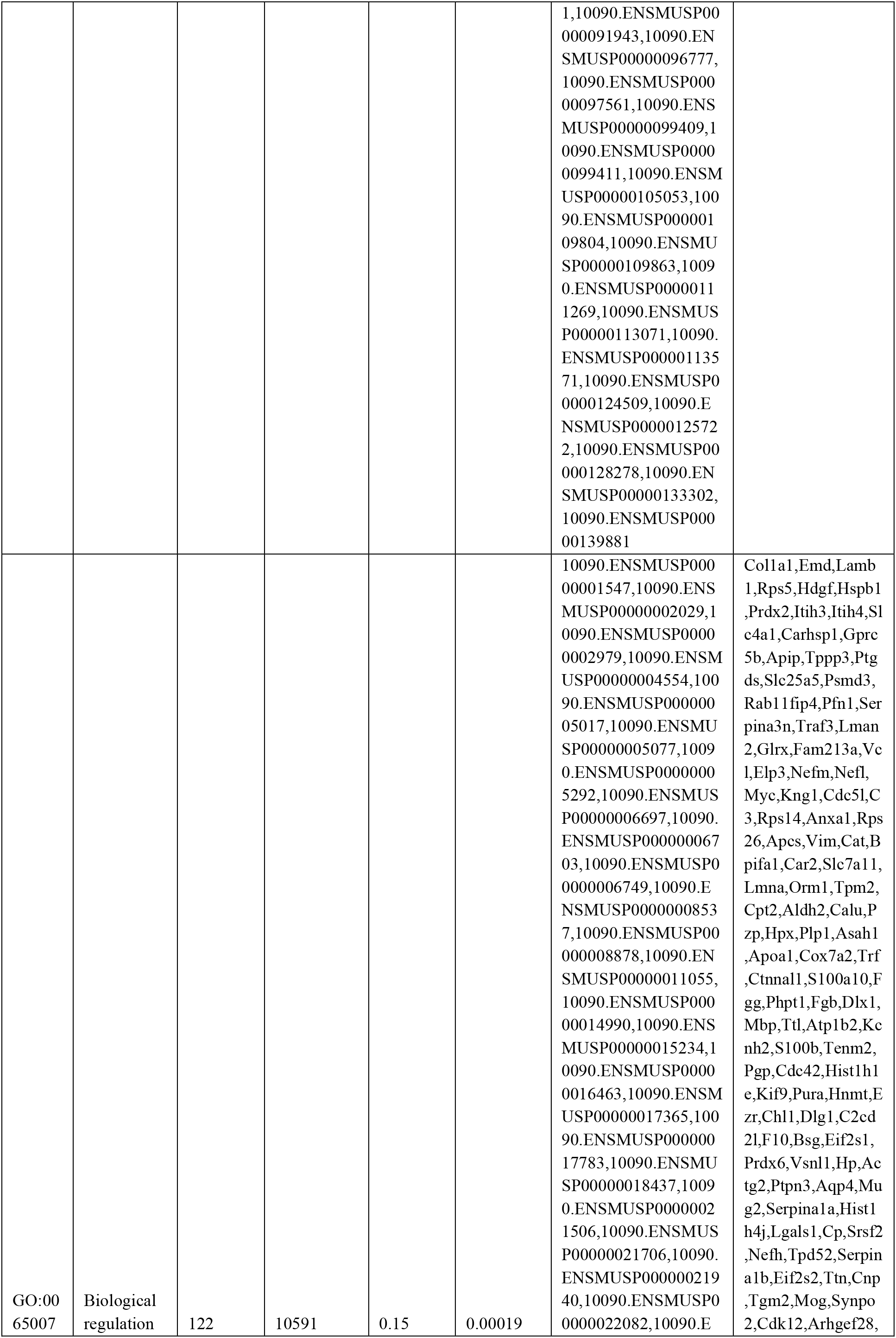
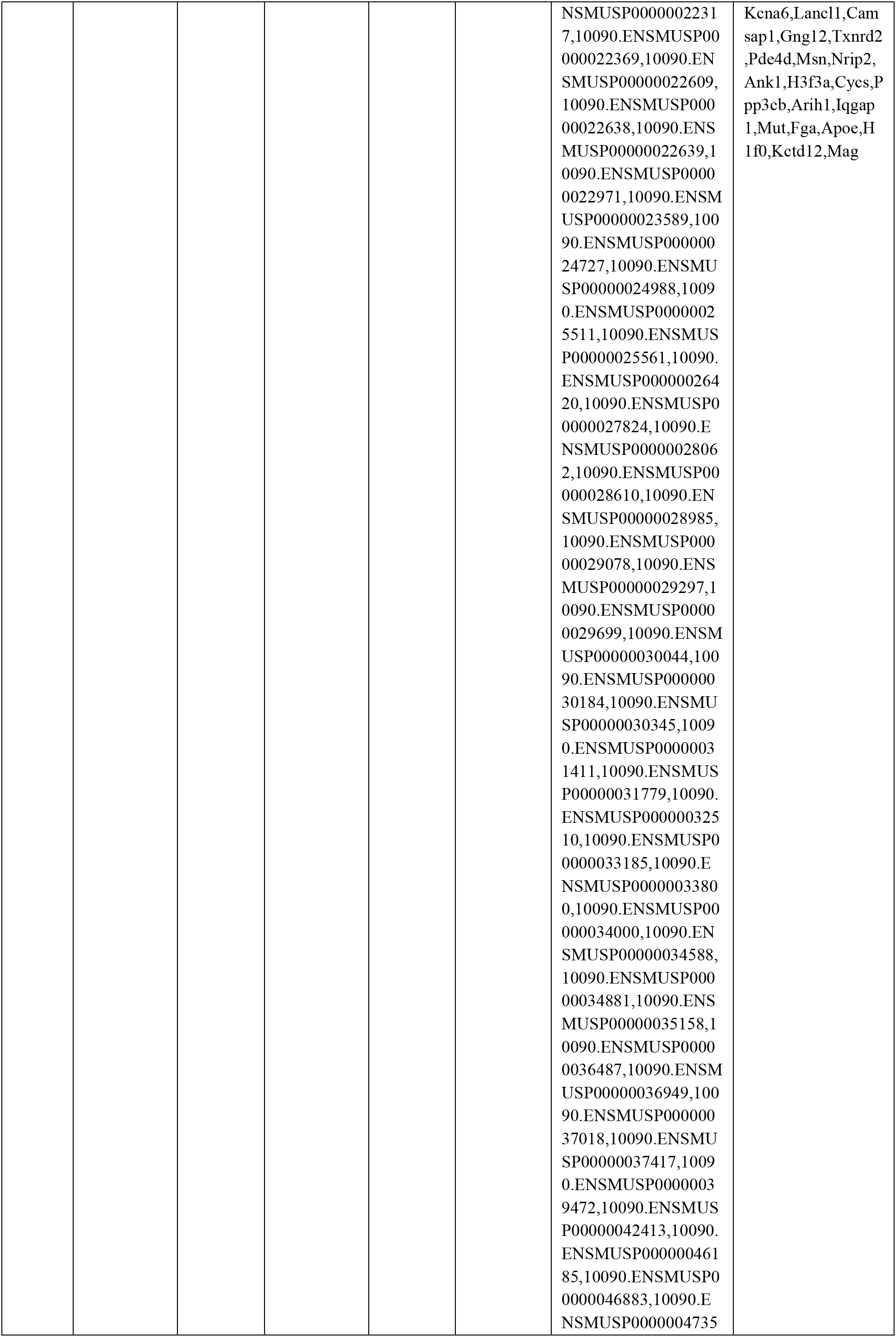
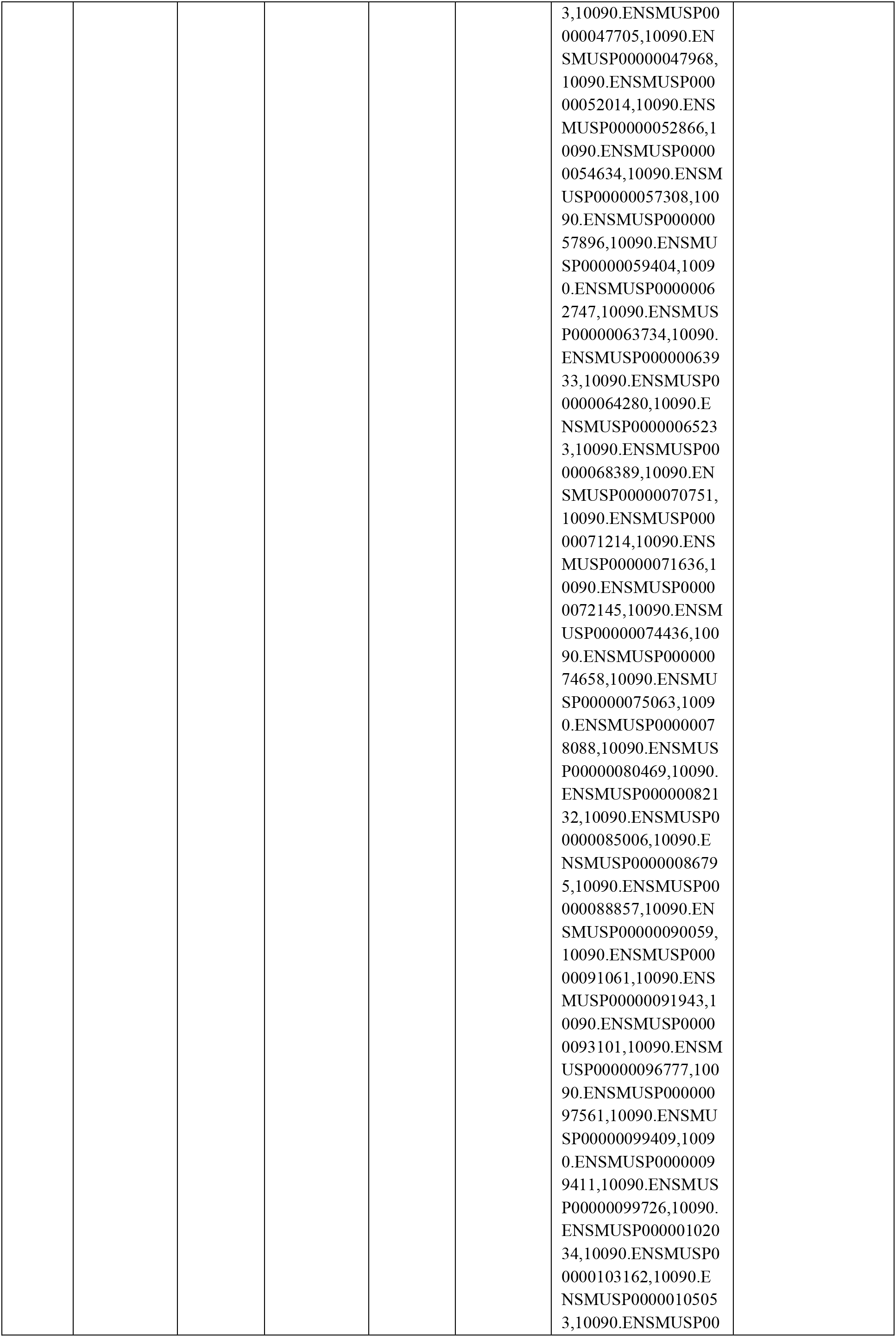
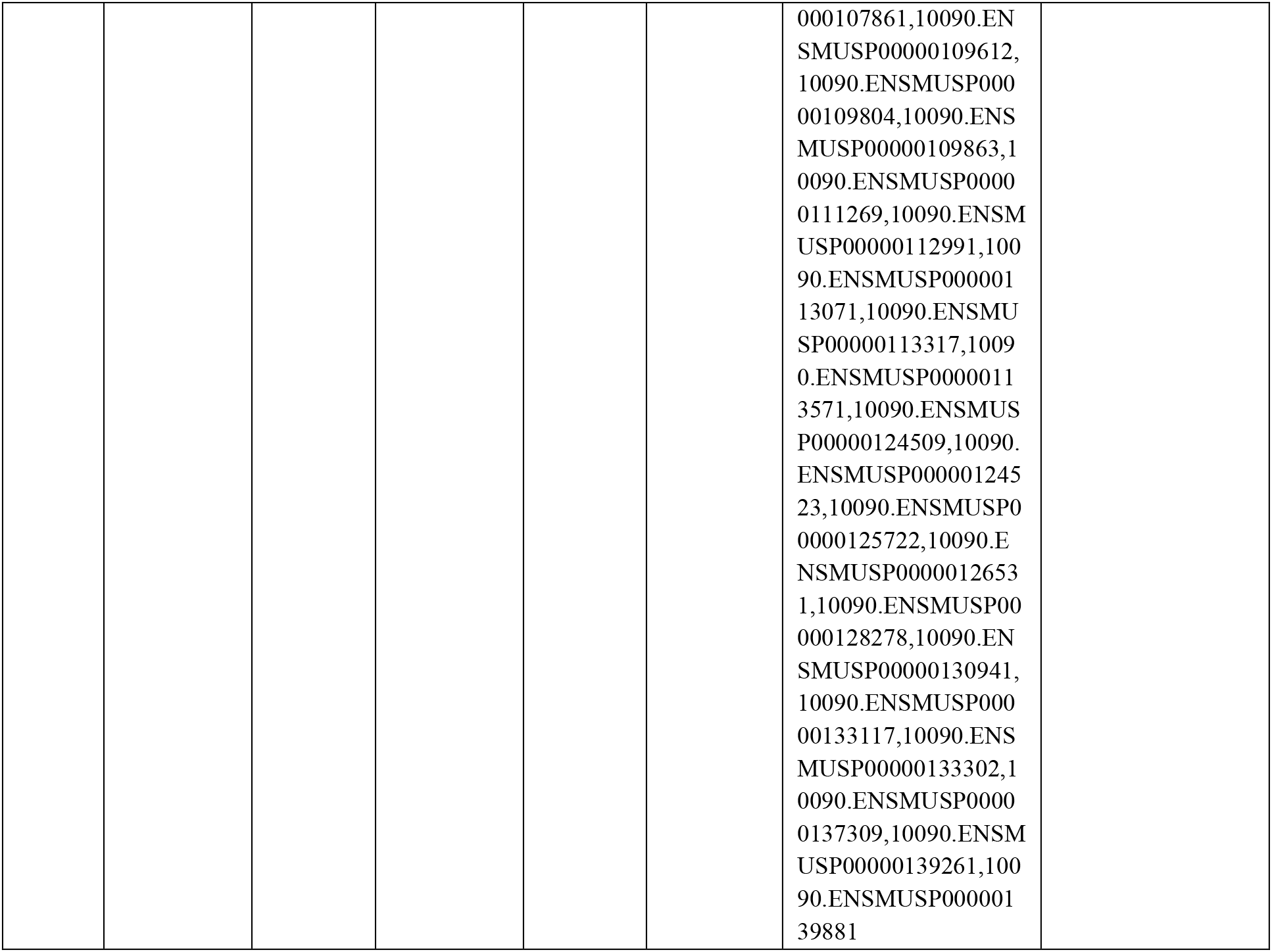
Biological Processes (GO) affected – Top 10 pathways

**Figure 2.**
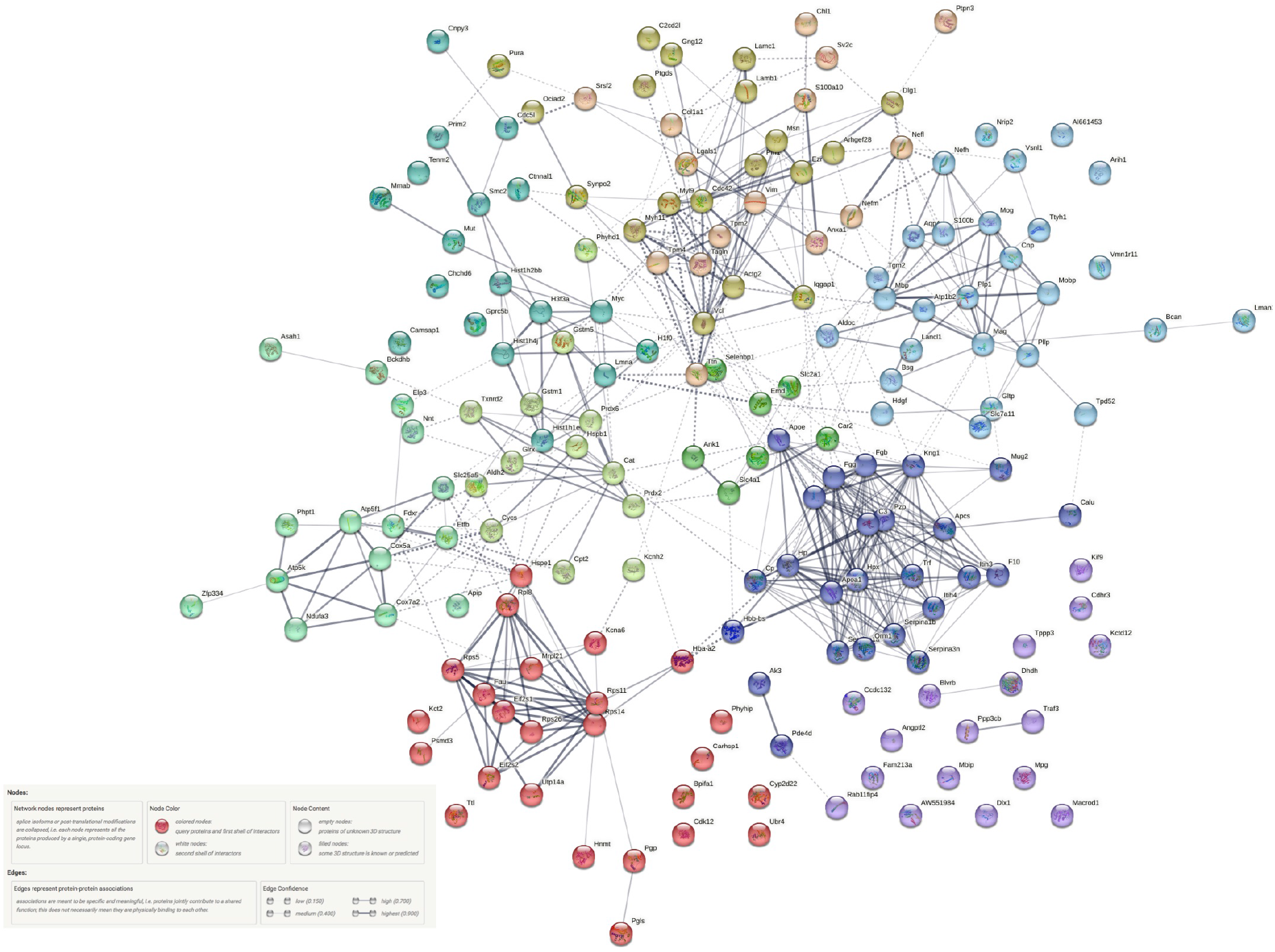
Medium confidence network map of upregulated proteins (Wt SD/Wt) - with thickness of lines depicting increasing confidence (0.15-0.90). k- means clustering with 10 clusters (dotted lines) was applied on these proteins. Both the network maps have been generated using String v.11.5.

Since stress might be affecting gene expression at the transcription level, we wanted to check whether upregulation was there at the transcript level of the genes whose protein levels were found upregulated in our study. Interestingly, we found equivalent changes in the transcript levels of *Ap1b1, Gat1, Hspe1, Nefm* and *Vim* (p<0.05; Fig 3).

**Figure 3.**
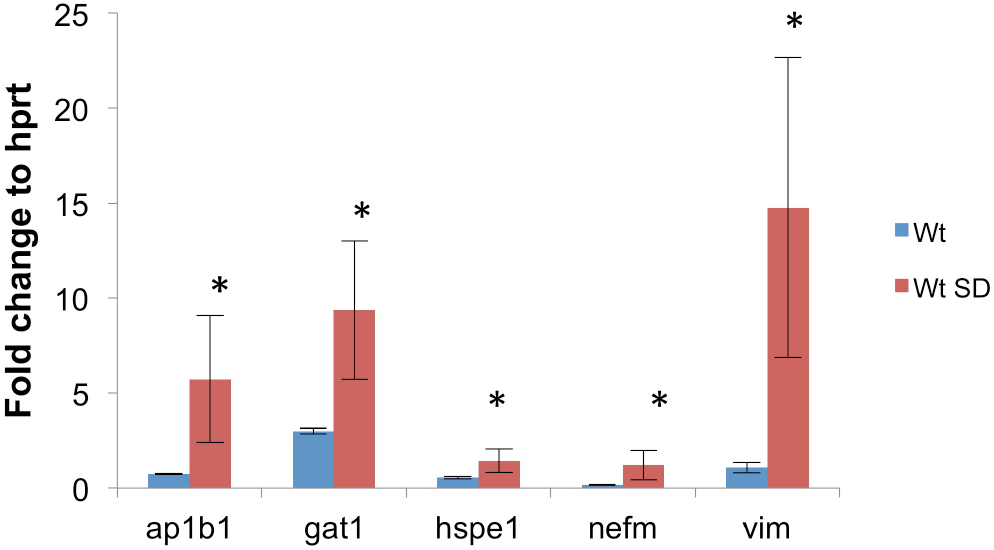
qPCR analysis of genes were carried out for proteins found upregulated. Equivalently transcripts of *Ap1b1, Gat1, Hspe1, Nefm* and *Vim* were found to be upregulated (p<0.05) in defeated group (Wt SD) as compared to unstressed controls (SD); indicating that the effect of social defeat stress might be at the transcriptional level.

## DISCUSSIONS

Social Defeat induces molecular as well as physiological changes in the brain. Multiple studies have been done to see the effect of chronic stress like social defeat in different brain regions like hippocampus, amygdala, pre-frontal cortex etc (Colyn *et al*, 2019; Fan *et al*, 2021; Iñiguez *et al*, 2014, 2016). In this present study, we analysed protein profile of hypothalamus, the major constituent of the HPA axis to understand the molecular changes that take place in hypothalamus on subjection to social defeat stress and whether we can correlate them to the response to stress.

A considerable fraction of the proteins found up regulated in the hypothalamus of stressed mice as compared to the unstressed controls belonged to the mitochondrial oxidation-reduction pathway and metabolism. Oxidative stress has been shown to be one of the major players behind pathogenesis of multiple brain disorders like Alzheimers, Parkinson, Huntington and Depression (Jiménez-Fernández *et al*, 2015; Liu *et al*, 2015). We found peroredoxin and thioredoxin, both potent anti-oxidants to be upregulated. We also found upregulation in Cytochrome c, which also has anti-oxidant properties. Cytochrome c can bind to super-oxides produced from electron transport chain in mitochondria and neutralise the free radical. Thus Cytochrome c aids in limiting and regulating the production of oxygen free radicals which may lead to oxidative stress (Bowman and Bren, 2008). Similarly there was an upregulation in the levels of glutathione S-transferases (GST). GST has been known for its role in inactivating electrophilic compounds and for de-toxification (Tirona and Pang, 1999; Townsend *et al*, 2003, 2005; Wu *et al*, 2004). Hence upregulation of these anti-oxidants might be a response to the build up of free radicals due to oxidative stress generated due to the chronic stress of social defeat in hypothalamus of mice.

We found upregulation in levels of neurofilaments NEFL, NEFM and NEFH. Transcript analysis indicates that NEFM is also transcriptionally upregulated. Neurofilaments are part of the axo-skeleton and also participate in axonal transport. NF may be accumulated in injury due to mechanical failure of the NF network (Meythaler *et al*, 2001). In multiple neurodegenerative conditions, NFs acts as a plasma (Ashton *et al*, 2021) and blood (Anderson *et al*, 2008) biomarkers for axonal injury and neuronal damage. Interestingly, NEFL has been found to be significantly upregulated in CSF of a concussed boxer even 30 days post-concussion indicating its usefulness as a biological marker for injury and recovery (Neselius *et al*, 2013). Taken together, the detection of elevated neurofilament in hypothalamus of mice brain post social defeat stress indicates two points. Firstly, it validates social defeat stress as a model for depression. Secondly, as in the case of concussed boxers, the present study uphold the potential of neurofilaments to be used as biomarkers in several human traumatic/stressful events akin to social defeat model in mice viz., bullying, aggression, subordination and humiliation. Recently few studies have shown that this might be just the case (Bacioglu *et al*, 2016; Chen *et al*, 2022; Travica *et al*, 2022).

In this study, we also found upregulation of Myelin Basic Protein (MBP) and associated proteins in the brain following stress in accordance with previous studies. MBP has been associated with neuronal death and trauma and has been found to be increased in CSF of dogs post traumatic brain injury (Bohnert *et al*, 2021; Su *et al*, 2012).

In our study we also found upregulation in the levels of several RPS subunits, like RPS5, 11,14, 26 (a ribosomal protein subunit). Ribosomal genes have been found to be upregulated in one more study that analysed de-regulated gene expression in the hypothalamus post social defeat stress (Smagin *et al*, 2016). This validates our finding at transcriptional level and also reflects upon the importance of protein translation under stress.

Changes were also found in levels of S100beta, a marker of astrocytes. An increase may indicate towards increased reactive astrocytes as a result of stress (Borella *et al*, 2003; Kleindienst *et al*, 2007). Similar lines hold for microglia, which has already been shown to be increased in social defeat model and in other stress models (Stein *et al*, 2017). An increase in CDC42 found in our proteomic dataset indicated to possible increase in microglial activation as a result of social defeat (Barcia *et al*, 2012; de Pablos *et al*, 2014).

We also found increased expression of Fibrinogen alpha, beta and gamma chains along with elevated levels of ApoA1. A complex of alpha-beta Fibrinogen can activate microglia thus leading to neuro-degeneration (Ryu *et al*, 2009). Fibrinogen and ApoA1 usually accumulate in the brain from the periphery (Ryu *et al*, 2009; Zhou *et al*, 2019) and their increased levels in hypothalamus might indicate compromised Blood Brain Barrier (BBB) as a result of social defeat stress. In fact recent investigations have shown that leaky BBB might be associated with different regions of brain in mice subjected to stressful paradigms including social defeat and associated with progression of depression like phenotype (Dion-Albert *et al*, 2022; Menard *et al*, 2017; Welcome and Mastorakis, 2020). This might also facilitate the movement of NEFM, NEFL and S100beta into the CSF – the elevated levels of which have been reported in clinical samples from individuals subjected to stress, particularly Traumatic Stress. This re-ignites the debate whether social defeat model is a model for depression like symptoms or ideally a model of PTSD (post traumatic stress disorder) (Richter-Levin *et al*, 2019).

Of the different proteins found up regulated, SLC4A1 also deserves special attention as it has been shown to be a promising biomarker in patients with MDD (Pajer *et al*, 2012).

Taken together, this present study gives information about de-regulated proteins in the hypothalamus of social defeat stressed mice. A large number of changes have been found in multiple studies in different brain regions. We found a considerable proportion of proteins in our data to be similar to those reported in the literature. Changes in hypothalamus are interesting bearing in mind that it interacts directly with the peripheral circulation with modified BBB. We report markers for neuronal death and atrophy and add weightage to the notion that neurofilaments like NEFL, NEFM and MBP may act as potential biomarkers in human stress and traumatic disorders.

## Supporting information

Proteins identified from both runs

List of differentially expressed proteins

Altered Biological processes (GO)

k-means clustering String analysis

## ACKNOWLEDGEMENT

We thank the Brain and Behavior Facility at CCMB for their help and support with the experiment. We thank the Proteomic Facility at CCMB for aiding in running the samples and analysis. We thank Dr. Satish Kumar, CCMB for his support and feedback.

## DATA AVAILABILITY

Data are available via ProteomeXchange with identifier PXD005644.

